# MICROVASCULAR ENDOTHELIAL CELL ADAPTATION TO HYPOXIA IS ORGAN-SPECIFIC AND CONDITIONED BY ENVIRONMENTAL OXYGEN

**DOI:** 10.1101/2020.08.25.265926

**Authors:** Moritz Reiterer, Amanda J Eakin, Aileen Burke, Randall S Johnson, Cristina M Branco

## Abstract

Microvascular endothelial cells (MVEC) are plastic, versatile and highly responsive cells, with morphological and functional aspects that uniquely match the tissues they supply. The response of these cells to oxygen oscillations is an essential aspect of tissue homeostasis, and is finely tuned to maintain organ function during physiological and metabolic challenges. Primary MVEC from two continuous capillary networks with distinct organ microenvironments, those of the lung and brain, were pre-conditioned at normal atmospheric (∼ 21 %) and physiological (5 and 10 %) O_2_ levels, and subsequently used to compare organ-specific MVEC hypoxia response. Brain MVEC preferentially stabilise HIF-2α in response to hypoxia, whereas lung MVEC primarily accumulate HIF-1α; however, this does not result in significant differences at the level of transcriptional activation of hypoxia-induced genes. Glycolytic activity is comparable between brain and lung endothelial cells, and is affected by oxygen pre-conditioning, while glucose uptake is not changed by oxygen pre-conditioning and is observed to be consistently higher in brain MVEC. Conversely, MVEC mitochondrial activity is organ-specific; brain MVEC maintain a higher relative mitochondrial spare capacity at 5% O_2_, but not following hyperoxic priming. If maintained at supra-physiological O_2_ levels, both MVEC fail to respond to hypoxia, and have severely compromised and delayed induction of the glycolytic shifts required for survival, an effect which is particularly pronounced in brain MVEC. Oxygen preconditioning also differentially shapes the composition of the mitochondrial electron transport chain (ETC) in the two MVEC populations. Lung MVEC primed at physioxia have lower levels of all ETC complexes compared to hyperoxia, an effect exacerbated by hypoxia. Conversely, brain MVEC expanded in physioxia display increased complex II (SDH) activity, which is further augmented during hypoxia. SDH activity in brain MVEC primed at 21 % O_2_ is ablated; upon hypoxia, this results in the accumulation of near-toxic levels of succinate in these cells. Our data suggests that, even though MVEC are primarily glycolytic, mitochondrial integrity in brain MVEC is essential for metabolic responses to hypoxia; these responses are compromised when cells are exposed to supra-physiological levels of oxygen. This work demonstrates that the study of MVEC in normal cell culture environments do not adequately represent physiological parameters found *in situ*, and show that the unique metabolism and function of organ-specific MVEC can be reprogrammed by external oxygen, significantly affecting the timing and degree of downstream responses.

**Graphical Abstract:** 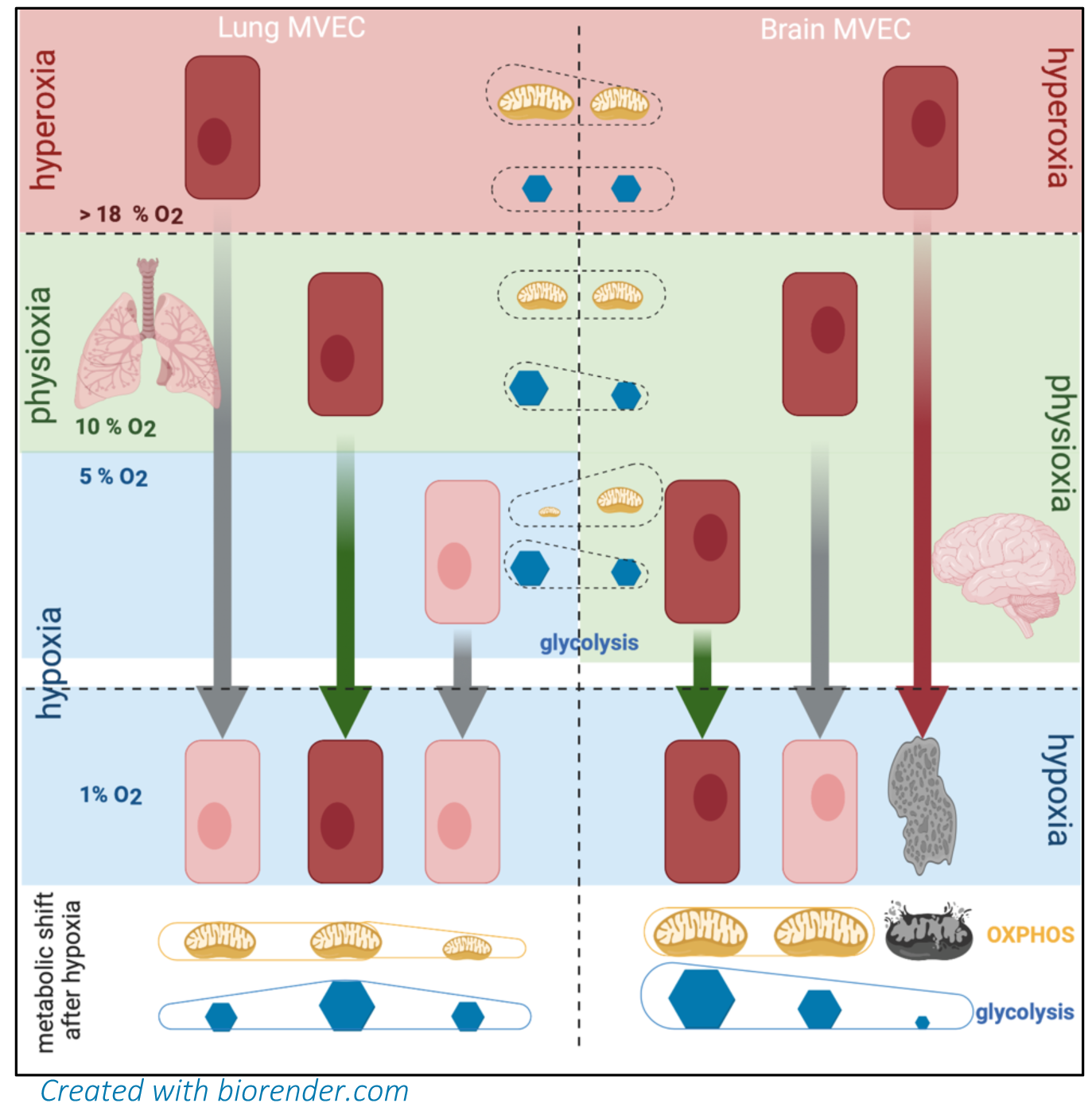

**In brief:** Hypoxia sensing by microvascular endothelial cells (MVEC) is organ-specific, and efficacy of response is affected by external oxygen. While glycolytic capacity is mostly maintained in MVEC regardless of organ or origin, mitochondrial function is required for adequate sensing and timely metabolic shift to glycolysis. Hyperoxygenation of MVEC compromises mitochondrial function, glycolytic shift and survival to hypoxia.

**Highlights:**

- Environmental O_2_ influences MVEC hypoxia response in an organ-specific fashion
- Brain MVEC are unable to respond and survive to hypoxia if hyperoxygenated prior to stress
- MVEC glycolytic capacity is not affected by O_2_, but the increase in glucose uptake and shift to glycolytic metabolism stifled and delayed in hyperoxidized MVEC
- High O_2_ ablates activity of mitochondria complex II in brain MVEC, significantly disturbing succinate levels Disruption of mitochondrial integrity compromises hypoxia sensing irrespective of glycolytic capacity

## Introduction

Microvascular endothelial cells (MVEC) are the most heterogeneous, versatile and specialised cells within the vascular tree (Augustin and Koh, 2017) (Aird, 2011), and are essential to organ function, which they regulate by locally adjusting nutrient, signalling and gas exchange rates to match organ demand to systemic availability (Reiterer and Branco, 2020). Angiocrine stimuli from resident MVEC mediate organ function and tissue microenvironment, which are also highly diverse in complex organisms (Zhang et al., 2020). Oxygen availability is dynamic and dependent on a tissue’s cellular components and metabolic activity, and as such, perception and response to oxygen levels by MVEC is critical to organ homeostasis, and effectively sidesteps autonomic control (Nolan et al., 2013; Reiterer and Branco, 2020). This plasticity underlies functions beyond perfusion and permeability, and ensures organ performance in a variety of physiological contexts, such as exercise or high altitude (Prefaut et al., 2000; Rojas-Camayo et al., 2018) or pathological conditions (Brahimi-Horn et al., 2007; Eltzschig and Carmeliet, 2011; Kent et al., 2011; Mistry et al., 2018), as well as during tissue remodelling and regeneration(Ding et al., 2010; Ding et al., 2011; Ding et al., 2015). MVEC are functionally and metabolically equipped to respond to hypoxia, and organisms are for the most part capable of seamlessly adjusting to O_2_ flux at systemic and local levels. When stretched too far or for too long beyond the normal physiological range, these cellular responses can escalate to exacerbate pathologies involving vascular dysfunction (Reiterer and Branco, 2019). Examples include diabetic retinopathy (Cai and Boulton, 2002) and atherosclerosis (Gimbrone and García-Cardeña, 2016)..

MVEC can also be exposed to supra-physiological oxygen levels (hyperoxia), often by elective use of oxygen for therapeutic purposes. Intra- and post-operative hyperoxia is frequently employed to increase neutrophil-mediated bactericidal activity, presumably by increased ROS production (Allegranzi et al., 2016; Hopf, 1997). Hyperbaric oxygen treatments are used for refractory diabetic wounds or to potentiate radiation efficacy (Chen et al., 2015; Goldman, 2009; Thom, 1989; Thom et al., 2011), as well as during diving (Banham, 2011). Unlike hypoxia, hyperoxia represents a non-physiological stimulus, which does not occur absent modern technology, and therefore, no evolutionary adaptation to it has taken place. Although clinically important, the effects of hyperoxia on MVEC function and plasticity have not been studied, with some exceptions on the effects on lung microvasculature (Ahmad et al., 2004; Kistler et al., 1967; Mach et al., 2011; Narula et al., 1998).

MVEC responses are presumed unique to the tissue they reside in, and in this study, MVEC from two continuous capillary networks from distinct microenvironments, lung and brain, were compared in their responses to hypoxia. We have found that environmental oxygen priming conditions the MVEC response to subsequent hypoxia. This conditioning affects both timing and amplitude of subsequent responses to hypoxia, and does so in an organ-specific fashion.

## Results

### Brain and lung microvascular endothelial cells respond differently to hypoxic stress

Primary murine brain and lung microvascular endothelial cells (lMVEC) were cultured in standard atmospheric conditions and subsequently exposed to hypoxia (1% O_2_) for up to 48 h. Cell viability was shown to decrease in both MVEC over time (Figure 1A), but the effect occurred earlier and was visibly more pronounced in bMVEC. Similarly, a real-time viability assay (Duellman et al., 2015) (Figure 1B) showed a more severely decreasing bMVEC growth curve.

**Figure 1:**
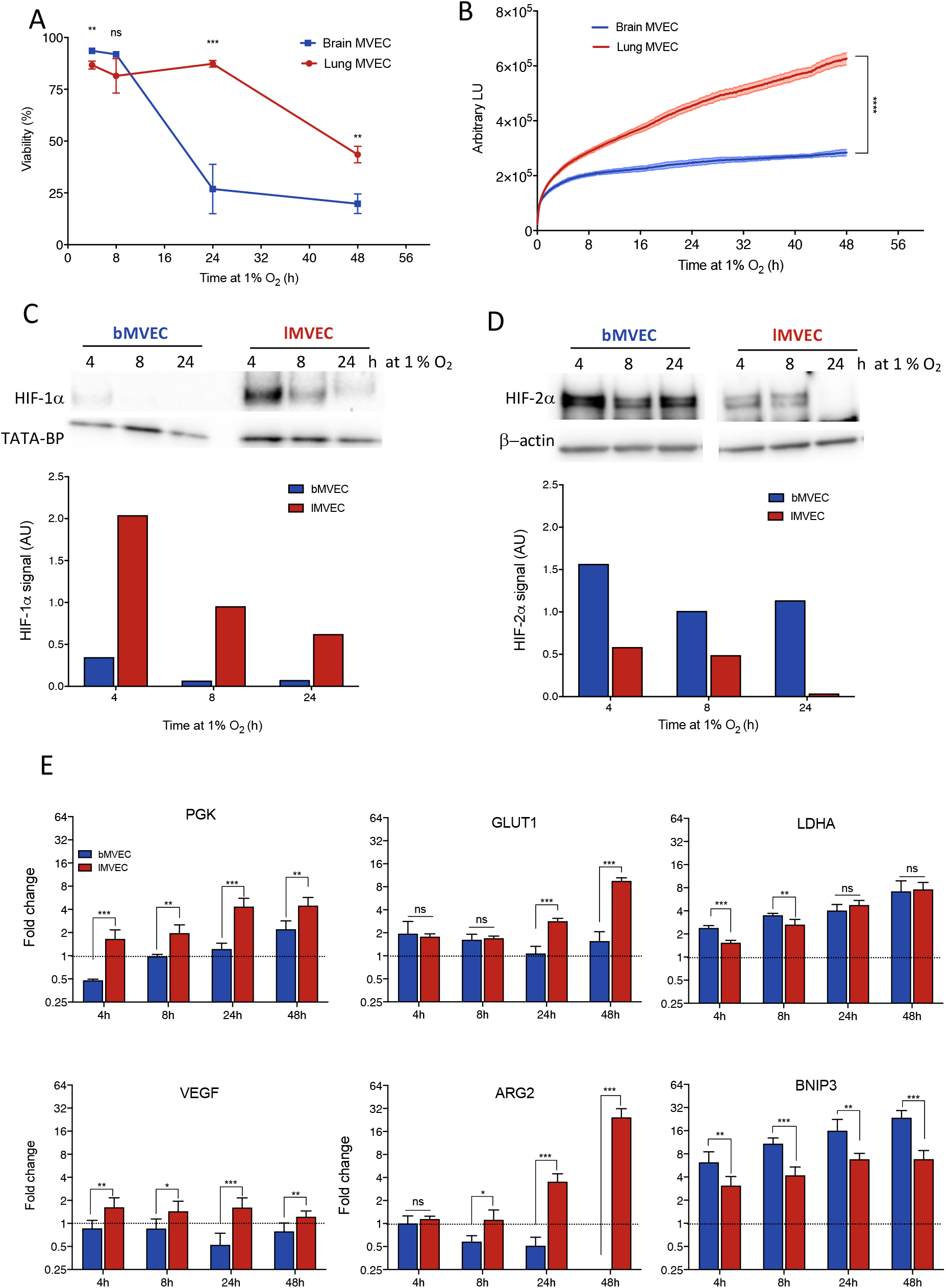
Lung and brain MVEC show organ-specific response and tolerance to hypoxia. (A) Viability of MVECs exposed to 1% O_2_ at t=0, measured by propidium iodide staining (n=3); t-tests corrected for multiple comparisons (Holm-Sidak) *p<0.05, **p<0.01, ***p<0.005) (B) Real-time viability of brain and lung MVECs exposed to 1% O2 at t=0, measured using the Real-Time Glo assay (Promega) (n=3). Shaded areas show SD, statistical differences were assessed by unpaired student’s t-test, using the area under the curve (****p<0.0001). (C,D) Western blot of HIF-1α and HIF-2α using nuclear extracts from brain and lung MVECs exposed to 1% O_2_ for the indicated amount of time. Quantification for this blot is shown, and was normalized to loading control (E) RT-qPCR for hypoxia targets PGK, VEGF, GLUT1, BNIP3, LDH-A, and HIF-2α target ARG2 in brain and lung MVEC, at 21 %, 10%, or 5% O_2_ baseline and after 1% O_2_. Data is shown as average fold-change ± SD (hypoxia/normoxia) (n=3). t-tests corrected for multiple comparisons (Holm-Sidak) using the log of fold change *p<0.05, **p<0.01, ***p<0.001)

As known key mediators of hypoxia response, activation of the two main HIF-α isoforms (HIF-1α and HIF-2α) was assessed in both MVEC populations, to investigate if discrepancies in the HIF-signalling pathway corresponded to differences in cell viability. HIF-1α expression showed the canonical transient upregulation in both lung and brain MVEC (Bartoszewski et al., 2019; Reiterer et al., 2020; Uchida et al., 2004), although consistently higher levels of this isoform was found in cells from the lung, at all time points (Figure 1C). HIF-2α protein levels decreased only at later time points for lMVEC, but bMVEC showed strikingly higher levels of HIF-2α protein throughout the hypoxia time course (Figure 1D).

Selected transcriptional HIF targets were quantified and, as expected, glycolytic genes PGK, LDH-A, and GLUT1 were upregulated in both MVEC, suggesting that HIF-α isoform preference does not necessarily underlie transcriptional activation of typical hypoxia induced genes; Even though the relative upregulation differed for each target, such as significantly more upregulated PGK transcript in lMVEC, or earlier upregulation of LDH-A in bMVEC, after 24 h of hypoxia the fold induction was no longer different between cell populations. One interesting distinction is that hypoxic lMVEC increasingly accumulated VEGF and ARG2 mRNA over time in hypoxia, but levels of either of those transcripts were seen to decrease in bMVEC, particularly unexpected for ARG2, considering this is considered primarily a HIF-2α target (Krotova et al., 2010) (Figure 1C). BNIP3, a HIF-1α target(Bellot et al., 2009; Keith et al., 2011; Liu and Frazier, 2015), was upregulated more strongly in brain than lMVEC, which correlates with the higher susceptibility to hypoxia-induced cell death seen in cells isolated from the brain (Figures 1A and B), but not HIF isoform preference.

These data show that MVEC from brain and lung display markedly distinct responses to hypoxia, including survival and HIF isoform stabilization, but the differences in hypoxia-driven transcriptional activation was not as strikingly affected.

*In vivo* oxygen tension in brain tissue (∼5 % O_2_) (Dings et al., 1998) is much lower than that found in the lung (∼10 %)(Wild et al., 2005), implying that bMVEC would be better equipped to survive an exposure to 1% O_2_. Paradoxically, bMVEC were more adversely affected by hypoxia, indicated by their reduced viability, and appeared unable to mount a suitable adaptive response when transferred to hypoxia, even though for many tissues, including the brain, 1 % O_2_ can still be considered physiological (Cater et al., 1961; Smith et al., 1977). Crucially, however, both organs contain much less oxygen than the O_2_ found in normobaric room air (Wenger et al., 2015), and thus the cellular responses observed at atmospheric oxygen tensions (∼ 21 % O_2_) are unlikely to represent physiological adaptations to hypoxia in the original tissue. To investigate this, primary MVEC were subsequently expanded in what was determined to be the best approximation to their typical physioxic state.

### Oxygen priming differentially conditions the hypoxia response of brain and lung MVEC

MVEC were cultured at either 10 % or 5 % O_2_ (physiological for lung and brain, respectively) in addition to standard 21 % O_2_, and hypoxia response evaluated for the same parameters as above. Upon transfer to 1% O2, both MVEC populations cultured in physioxia remined more viable than those primed at 21 % O_2_ (Figure 2A). HIF-α isoform levels were again assessed after 4 h of hypoxia and, as before, HIF-1α signal was consistently higher in lMVEC than bMVEC, and this was significant when cells were expanded at 21 % or 10 % O_2_ (but not 5 %) prior to hypoxia. In both cell populations, HIF-1α levels after hypoxia were highest if MVEC were cultured at 10% O_2_ irrespective of their tissue of origin (Figure 2B); The quantification of HIF-1α is shown in Figure 2C, where each data point refers to one biological replicate and dashed lines link lung and brain MVEC from the same experiment (and quantified from the same gel). Conversely, and similarly to what was shown in Figure 1D, HIF-2α protein levels were higher in hypoxic bMVEC compared to lMVEC (Figure 2D); Quantification of hypoxic HIF-2α signal is plotted in Figure 2E and, as above, signals from same experiment are linked by a dashed line. Significant differences between MVEC populations were only seen at the two highest O_2_ concentrations.

**Figure 2:**
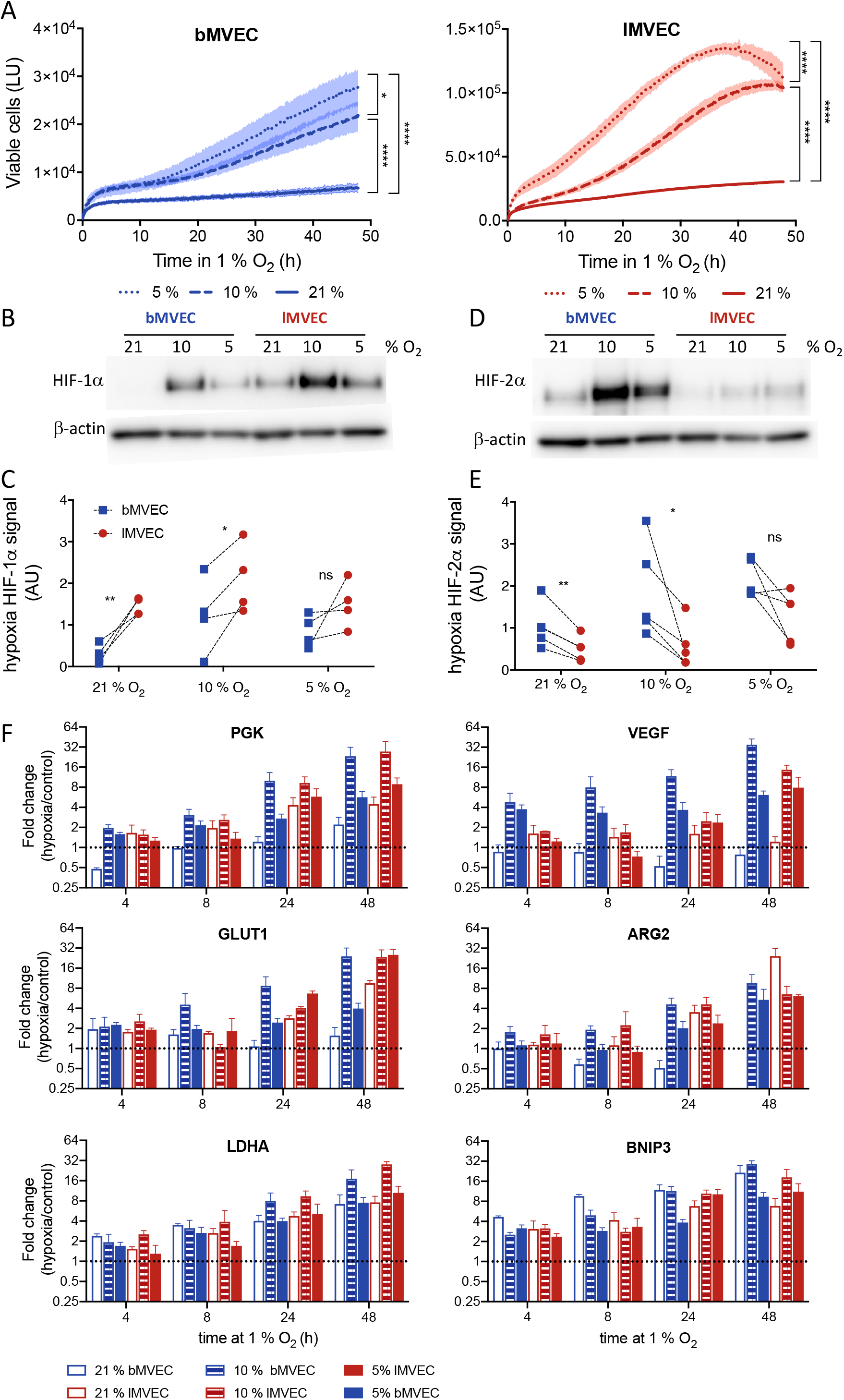
MVEC response to hypoxia is organ-specific and dependent on O2 priming. (A) Real-time viability of brain (left, blue) and lung MVEC (right, red) maintained at 21 %, 10%, or 5% O_2_ and exposed transferred to hypoxia (1% O_2_) at t=0 was measured using a Real-Time Glo assay (Promega) (n=3). Shaded areas show SD, statistical differences were assessed by 2-way ANOVA with Holm-Sidak’s multiple comparison test, using the area under the curve (*p<0.05, ****p<0.0001). (B) Representative western blot of HIF-1α using nuclear extracts from brain and lung MVEC exposed to 1% O_2_ for 4 h, and previously expanded at 21 %, 10 %, or 5 % O_2_ (C) Quantification of HIF-1α signal obtained by densitometry and normalized to the loading control; the lines connect samples from the same experiment (n=4; paired t-test, *p<0.05, **p<0.01) (D) Representative western blot of HIF-2α using nuclear extracts from brain and lung MVEC exposed to 1% O_2_ for 4 h, and previously expanded at 21 %, 10 %, or 5 % O_2_(E) Quantification of HIF-2α signal obtained by densitometry and normalized to the loading control; the lines connect samples from the same experiment (n=4; paired t-test, *p<0.05, **p<0.01) (F) RT-qPCR for hypoxia targets PGK, VEGF, GLUT1, BNIP3, LDH-A, and HIF-2α target ARG2 in brain and lung MVEC, at 21 %, 10%, or 5% O_2_ baseline and after 1% O_2_. Data is shown as average fold-change ± SD (hypoxia/baseline at 4h) (n=3). Statistical differences were assessed by 2-way ANOVA with Holm-Sidak’s multiple comparison test, using the log of fold change; the results of the statistical analysis are listed in Table S1.

To investigate if these results reflected a change from the baseline HIF-α levels or were consistently high or low in specific MVEC populations, HIF protein levels before hypoxia were also quantified (Supplementary Figure S1A). Both isoforms were stable at physiological oxygen, underscoring their role in normal EC function; lMVEC has higher levels of HIF-1α, whereas HIF-2α at baseline was low in both MVEC. Additionally, and to allow a direct assessment of HIF-α change of signal following hypoxia, hypoxic and physioxic nuclear protein extracts were run on the same gel (Supplementary Figure S1B, C), confirming that hypoxia induction of HIF-1α occurs primarily in lMVEC, except when they are expanded at 5 % O_2_, indicating that this condition is sub-physiological (or even hypoxic) for lMVEC. Contrary to the association of HIF-1α activation with the early hypoxia response, 4 h of hypoxia led to preferential induction of HIF-2α in bMVEC at physiological baseline O_2_, and to much higher levels than lMVEC induce HIF-1α. This is likely a result of elevated HIF-1α levels in physioxic atmospheres. HIF-2α induction in bMVEC was higher when cells were primed at lower O_2_, whereas HIF-1α induction in lMVEC was lower. Conversely, HIF-1α induction in bMVEC and HIF-2α induction in lMVEC seem independent of baseline O_2_ (Supplementary Figure S1B, C).

Transcript levels of HIF targets are consistently more strongly upregulated in cells maintained in physiological O_2_ (Figure 2F, Supplementary Table S1). Most strikingly, VEGF and ARG2 mRNA levels were induced by hypoxia in bMVEC only if the cells had been cultured at 10% or 5% O_2_. The autophagy marker BNIP3, however, was induced most strongly in bMVEC cultured at 21 % O_2_, correlating with their reduced survival to hypoxia in this condition. To extricate if the mRNA fold-change was skewed by transcript levels at baseline, the relative abundance of hypoxia-responsive transcripts before hypoxia treatment was plotted (Supplementary Figure S2). Indeed, some intrinsic differences are seen in the relative abundance of individual transcripts between the two MVEC populations, such as lower levels of LDH-A and VEGF in bMVEC. Baseline transcript levels were also different depending on O_2_ priming, and much higher GLUT1 levels in lMVEC maintained at 10 % O_2_. Overall, the largest increases in transcript levels of hypoxia response genes were generally seen in cells grown in physiological oxygen, especially at 10 % O_2,_ and mostly for bMVEC.

To investigate if discrepancies were due to upstream regulation of HIF stabilisation (prolyl hydroxylases PHD1-3) or transcriptional activity (FIH) (Mahon et al., 2001; Lando, Peet, Gorman, et al., 2002), relevant protein levels were investigated by western blot (Supplementary Figure S3A). Protein levels of the canonical regulators of HIF were quantified, but no O_2_-dependent patterns or tissue-specific differences were seen to significantly differ between MVEC (Supplementary Figure S3A, S3B). PHD2 levels were slightly reduced in bMVEC at 10 % and 5% O_2_, and PHD3 was consistently higher in those cells, in all O_2_ conditions, compared to lMVEC. FIH expression was significantly elevated in MVEC maintained in 21 % O_2_, irrespective of tissue of origin. While these results may not explain the HIF isoform levels, it may contribute to the dampened hypoxia response of MVEC cultured in hyperoxia. The levels of HIF-α mRNA were also measured (Supplementary Figure S2B), and almost entirely reflect the baseline protein abundance quantified by WB, despite the traditional assumption that HIF is mostly post-translationally regulated. However, lMVEC grown at 21 % O_2_ displayed much lower levels of HIF-1α mRNA than any other sample, a trend not reflected at the protein level.

The data above shows that MVEC viability is affected by oxygen priming, yet even though HIF-α isoform activation is tissue-specific, it does not appear to affect HIF-dependent transcriptional activation of hypoxia-related targets, or viability when cells are primed in physioxia, and as such the different tolerance seen between lung and brain MVEC appear largely independent from HIF stabilization.

### Glycolytic activity of MVEC is altered as a result of oxygen priming

Endothelial metabolism has been shown to underlie endothelial function, and EC are widely believed to be consistently and intrinsically glycolytic (Krützfeldt et al., 1990). However, a further shift towards glycolysis is key to enable adaptation and survival to hypoxic conditions. Glycolytic stress tests were performed to compare glycolytic metabolism in MVEC from brain and lung tissues, using extracellular acidification rate (ECAR) as a readout of glycolytic activity at baseline and upon complete mitochondrial inhibition with Oligomycin (maximal capacity) (Teslaa and Teitell, 2014). Representative graphs are shown in Figure 3A, and summaries of average glycolytic parameters quantified in the two MVEC populations are presented in Figure 3B. Baseline glycolysis, as expected, was higher in cells maintained in environments with lower O_2_. Although maximal glycolytic capacity (and thus spare capacity) was comparable between the two MVEC at all O_2_ tensions, lMVEC had slightly but significantly higher basal glycolytic activity at physiological O_2_ tensions than bMVEC (Figure 3B). Changes in glycolytic function following adaptation to hypoxia were subsequently assessed (Figure 3C, D). Here, cells expanded in the three different oxygen environments were transferred to 1% O_2_ for 24 h prior to the assay, to stabilise any metabolic changes resulting from adaptation to hypoxia. Cell viability was confirmed and cell density optimised for each MVEC population (see methods), to avoid assaying non-viable cells or induce anoxia in assay wells. Adaptation to hypoxia in terms of glycolytic activity is both oxygen-dependent and organ-specific (Figure 3C). lMVEC maintained an inverse correlation between O_2_ levels and glycolytic rates seen at baseline (Figure 3B); following adaptation to hypoxia, lMVEC showed a basal glycolytic rate lower than that of bMVEC if the cells had been grown at 21 %, similar if the cells had been grown at 10%, and higher if the cells had been grown at 5 % O_2_ (Figure 3D). Interestingly, hypoxic bMVEC had a larger glycolytic spare capacity following 24 h of hypoxia in all condittions (significant at higher O_2_ levels). Indeed, bMVEC are much more likely to encounter such low levels of oxygen *in vivo*, and as such, by maintaining glycolytic plasticity, would be better equipped to adjust to low O_2_. This, however, is in stark contrast with the fact that bMVEC are more susceptible to hypoxia, albeit only when primed at 21 % O_2_ (Figures 1, 2).

**Figure 3:**
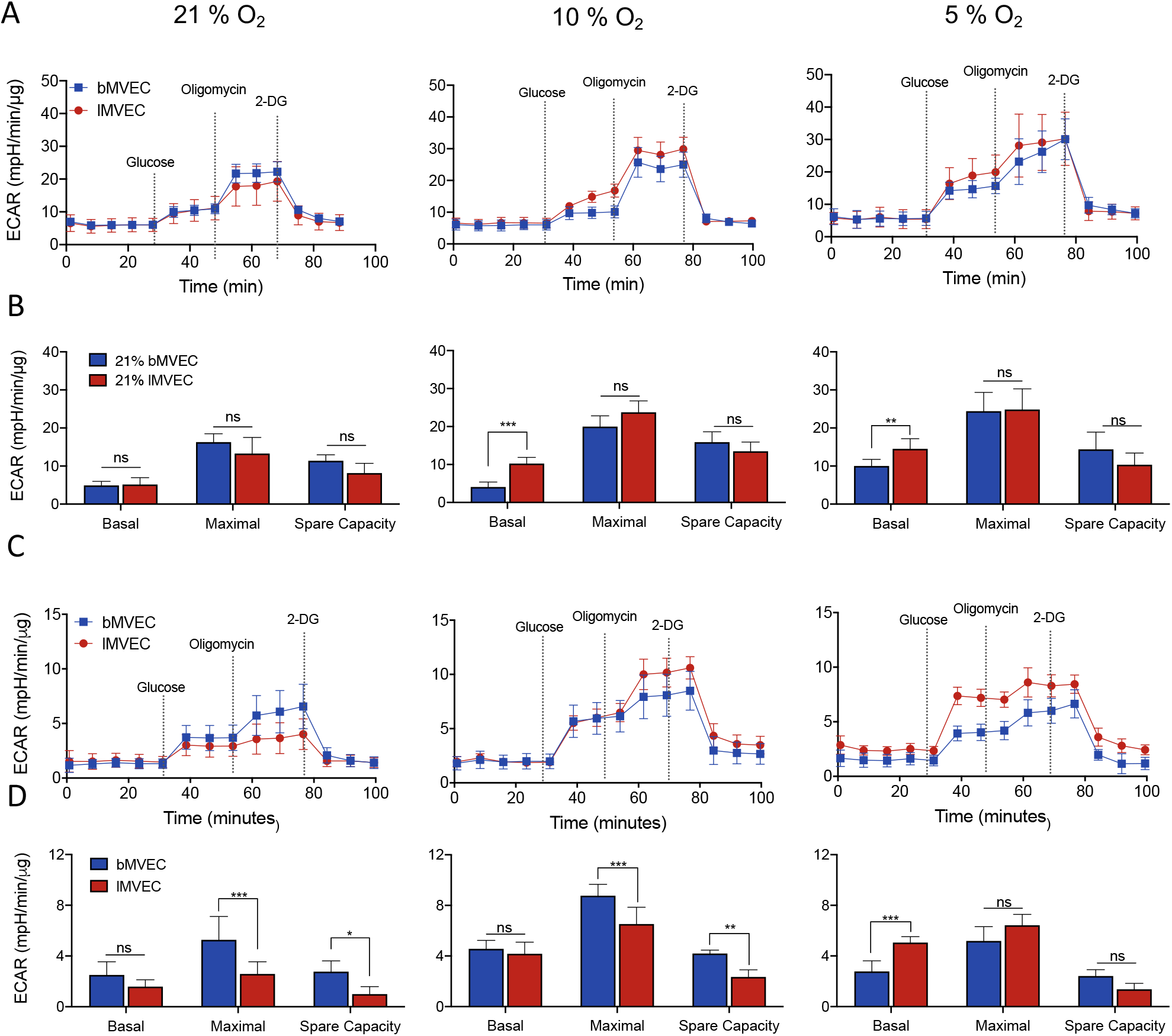
MVEC glycolytic activity is organ-specific and affected by O2 priming. (A) Representative charts of baseline glycolytic stress tests of brain and lung MVEC cultured at 21 % O_2_ (left), 10 % O_2_ (middle), or 5 % O_2_ (right). (B) Glycolytic parameters calculated from each curve from (A) are shown as average ± SD (n ≥ 4). Statistical analysis was done using 2-way ANOVA with Holm-Sidak’s multiple comparison test (*p<0.05,**p<0.01, ***<0.001) (C) Representative charts of glycolytic stress tests of brain and lung MVEC after 24 h of hypoxia (1 % O_2_), from cells primed at different O_2_; assays were carried out at 1% O_2_ (D) Glycolytic parameters calculated from each curve from (C) are shown as average ± SD (n ≥ 4). Statistical analysis was done using 2-way ANOVA with Holm-Sidak’s multiple comparison test (*p<0.05,**p<0.01, ***<0.001)

To further investigate this conundrum, ECAR was measured immediately upon transfer to 1 % O_2_, such that the glycolytic shift could be observed in real-time, instead of allowing the cells 24 h to adjust. This revealed that oxygen priming drastically affects MVEC adaptation to hypoxia (Figure 4A), both in terms of timing and amplitude (note y-axis matched scale). Both MVEC grown in 21 % O_2_ (left) showed a very mild increase in ECAR, which was further delayed in bMVEC. If primed at 10 % O_2_ (middle), lMVEC increased glycolytic activity within the first 5 h, to more than 2-fold than the same cell population expanded at 21 % O_2_. For the same conditions (21% and 10%), increase in ECAR for bMVEC upon hypoxia exposure was still very mild. However, when MVEC were maintained at 5 % O_2_ prior to hypoxia (right), the ECAR measured in bMVEC increased within minutes, to an extraordinary extent, highest seen for either cell type in any condition. This experiment demonstrates that bMVEC are significantly more responsive to hypoxia, but only if kept in physioxia. On the other hand, lMVEC moved from 5 % to 1 % O_2_ are unable to further increase ECAR, corroborating that 5 % O_2_ is already seen as hypoxia by these cells.

**Figure 4:**
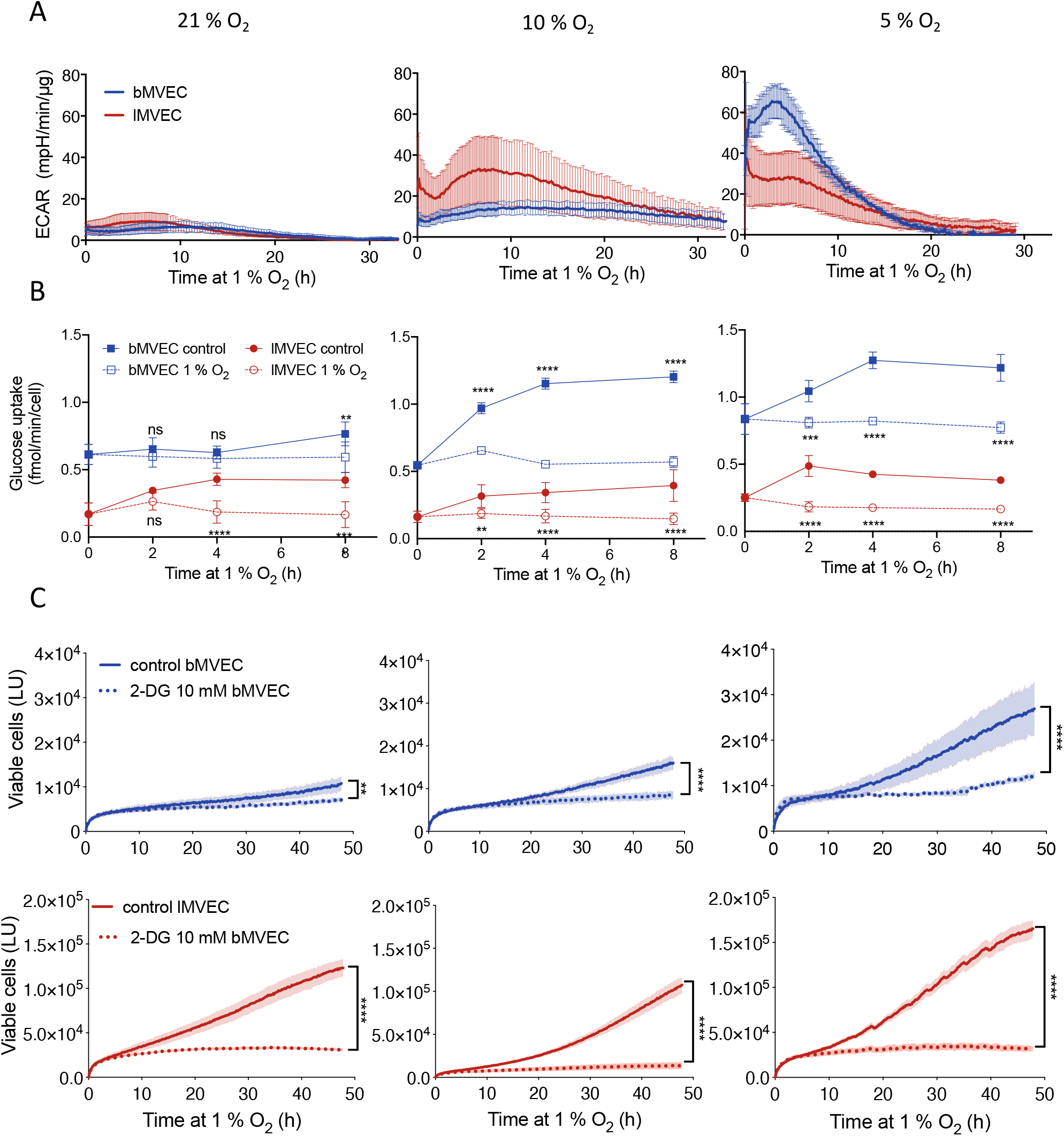
Timing and amplitude of metabolic shift to glycolysis is organ-specific and oxygen-dependent. (A) Representative charts of real-time measurement of ECAR changes upon transfer to 1% O_2_ at t=0, in lung and brain MVEC previously expanded at different O_2_ levels (21% O_2_, left), 10% O_2,_ middle, or 5% O_2_, right) (n≥ 4) (B) Glucose uptake in MVEC expanded at different O_2_ levels at baseline (dashed line) or under hypoxia (solid line), was measured at the indicated times using the Glucose Uptake-Glo assay (Promega) Data shown as average ± SD, and statistical differences were assessed by 2-way ANOVA with Holm-Sidak’s multiple comparison test (**p<0.01, ***<0.001, ****p<0.0001), n=3. (C) Real-time viability of brain (top, blue) and lung (bottom, red) was assessed every 10 min using a Real-Time Glo assay (Promega) in MVEC transferred to 1% O_2_ at t=0 and treated with 10 mM 2-deoxy glucose (dotted line) or PBS vehicle control (solid line). N=6 for each condition. Shaded areas show SD, statistical differences were assessed by unpaired student’s t-test, corrected for multiple comparisons (Holm-Sidak) using the area under the curve (**p<0.01, ****p<0.0001).

Glucose uptake assays were also performed in real time (Figure 4B). This experiment revealed that bMVEC have intrinsically higher rates of glucose uptake than lMVEC, irrespective of O_2_ availability (Figure 4B, Supplementary Figure S4A), or indeed GLUT1 transcript levels (Figure 1E, 2F, Supplementary Figure S2). As expected, glucose uptake increases in all MVEC upon transfer to hypoxia (Figure 4B), except in bMVEC primed at 21 % O_2_ (Figure 4B, left), which appear inept at increasing glucose uptake until 8 h after hypoxia exposure. Glucose uptake is an essential step in triggering a concomitant increase in glycolytic activity, and is in agreement with the data Figure 4A (left), showing bMVEC have a delayed perception of O_2_ levels when maintained in 21 % O_2_.

Viability assays performed in the presence of 10 mM 2-DG, a competitive inhibitor of glycolysis were performed to investigate if the increased hypoxic viability of MVEC grown at lower oxygen levels was indeed due to dependence on glycolytic metabolism. Removing the ability to use glucose for cellular energetic demands under hypoxia resulted in arrest of proliferation, irrespective of the glycolytic reserve of each MVEC population or culture condition (Figure 4C).

Combined, the results shown above strongly suggest that high oxygen levels impair MVEC perception of, and therefore response to, a hypoxic challenge, and this is more striking in MVEC that originate from brain tissue.

### Effects of oxygen priming on mitochondrial respiration

Following the assessment of glycolytic capacity and response in different MVEC populations, the effect on mitochondrial metabolism was investigated and compared as a function of oxygen availability. Oxygen consumption rates (OCR) were calculated from mitochondrial stress tests. Mitochondrial basal and maximal respiration rates, predictably, directly correlated with oxygen levels in both MVEC: those grown at 21 % O_2_ have the highest OCR, whereas those maintained at 5% O_2_ were the least reliant on mitochondrial respiration, as their metabolic preference is, naturally, glycolysis (Figure 5A, note mismatched y-axis scales, and Figure 5B). This pattern was considerably more pronounced in lMVEC, and as a result the maximal OCR of lMVEC grown at 21 % O_2_ was higher than that of bMVEC (Figure 5A,B left), whereas at 5% O_2_ that pattern was strikingly reversed (Figure 5A, B, right), and bMVEC retained a proportionally higher mitochondrial reserve. At 10 % O_2_ (Figure 5A,B, middle panels) there was no difference between the two MVEC populations. Thus, at physiological O_2,_ bMVEC maintain higher relative mitochondrial spare capacity than their lung counterparts. Mitochondrial respiration parameters were assessed also following 24 h at 1 % O_2_. Similarly to what was seen at baseline (prior to hypoxia), basal respiration rates were identical between the two MVEC populations primed at 21 % O_2_ (Figure 5C, D, left). Maximal respiration was seen to decrease only in lMVEC, whereas bMVEC grown in physiological O_2_ retained a significantly higher maximal respiration rate compared to lMVEC (Figure 5C, D, middle and right panels).

**Figure 5:**
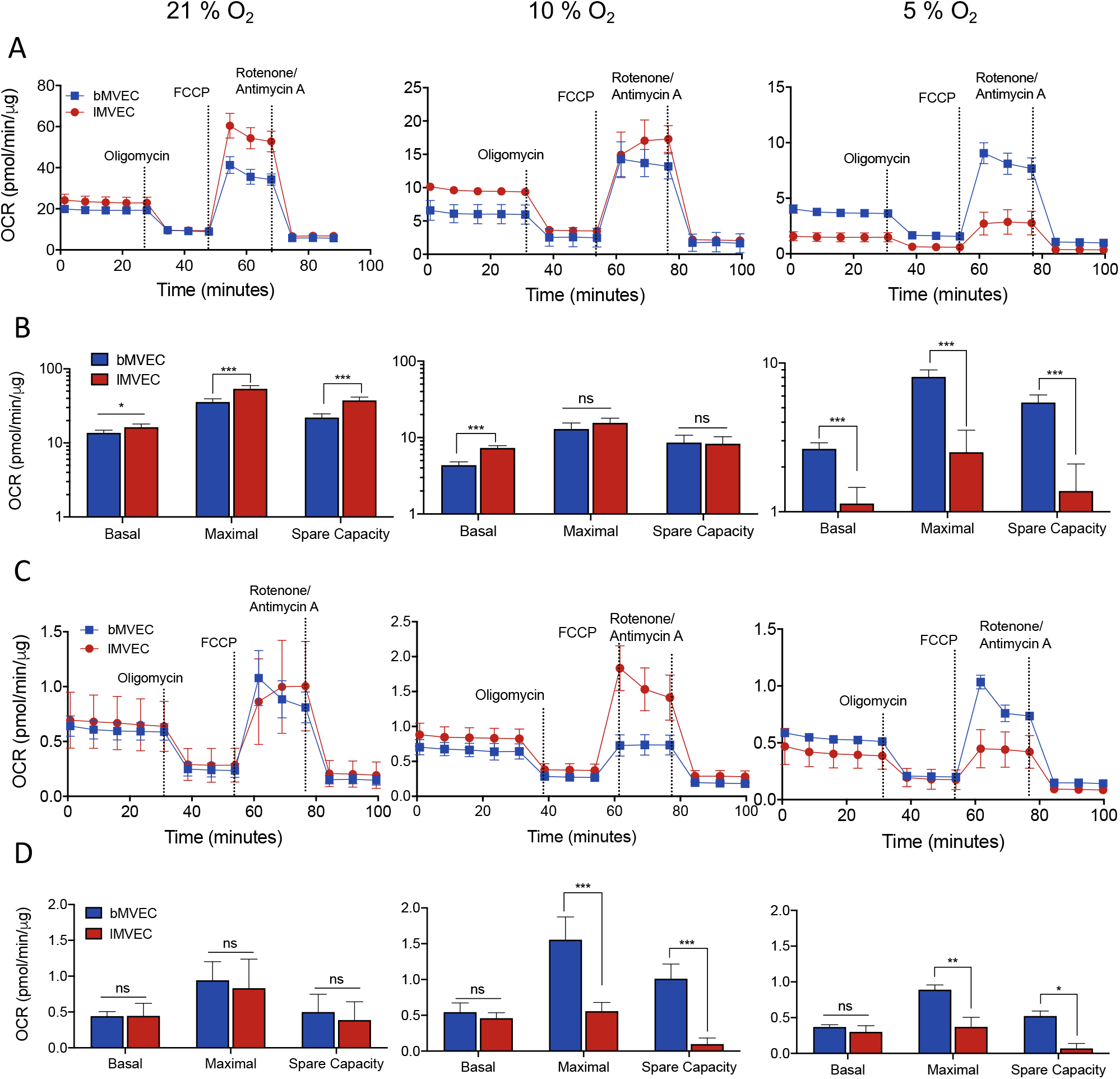
Mitochondrial metabolism of MVEC at baseline and after hypoxia is conditioned by oxygen priming and organ of origin. (A) Representative charts of mitochondrial function at baseline, in MVEC cultured at different O_2_ (21 %, left, 10 % O_2_ middle, or 5 % O_2_ right) (B) Mitochondrial metabolic parameters were calculated from each curve from (A). Data are shown as average ± SD (n≥ 4), statistical analysis was done using multiple t-tests with Holm-Sidak’s multiple comparison test (*p<0.05,**p<0.01, ***<0.001) (C) Representative charts of mitochondrial function after adaptation to hypoxia (1 % O_2_ for 24 h prior to start of the assay). Assays were carried out at 1% O_2_ (D) Mitochondrial metabolic parameters of assays carried out after hypoxia were calculated from each curve from (C). Data are shown as average ± SD (n ≥ 4), Significance assessed with 2-way ANOVA with Holm-Sidak’s multiple comparison test (*p<0.05,**p<0.01, ***<0.001)

These data indicate that bMVEC have wider metabolic plasticity, and overall higher mitochondrial activity than their lung counterparts. However, it is also apparent that bMVEC are more vulnerable to high oxygen levels. To confirm that the EC metabolic profile is reflected in functional aspects of MVEC behaviour, a migration assay was performed to assess alterations in EC function as a result of O_2_ priming (Supplementary Figure 4B, C). This was done in the absence (Supplementary Figure 4B) and presence of the proliferation inhibitor Mitomycin C (MM) (Chen et al., 2013) (Supplementary Figure 4C,D), thus discriminating closure due to proliferation and migration, or migration alone, respectively. MVEC maintained in their respective physioxia closed the scratch wound more effectively than in either of the alternative atmospheres, and were slowest to migrate when in hyperoxia. bMVEC were generally less motile than those from lung, and the effect of O_2_ pre-conditioning was less pronounced; also, and seeing that the fastest closure occurred in cells at 5% O_2_ suggests that bMVEC are intrinsically less migratory, and rely more on cell division for wound closure than do lMVEC, which appear generally less susceptible to the presence of MM.

### Effects of O_2_ on mitochondrial ETC complexes is organ-specific

Having investigated the effects of varying oxygen levels on mitochondrial metabolism, the impact on the mitochondrial ETC composition was further investigated. Total protein was isolated from bMVEC and lMVEC at baseline and following 24 h of adaptation to 1 % O_2_, for each oxygen condition. These were resolved by SDS-PAGE, probed with a mitochondrial ETC antibody cocktail (Sim et al., 2018) and quantified upon normalization against a positive control (Figure 6). A representative blot of lMVEC protein (Figure 6A) and the ratio of signal for each complex was compared to that seen at 21 % O_2_ (Figure 6B). In lMVEC, there was a positive (and expected) correlation between oxygen levels and mitochondrial complex protein levels; cells grown at lower O_2_ had lower levels of mitochondrial ETC complexes than cells grown at 21 % O_2_. This was further (and in most cases, significantly) exacerbated in MVEC following adaptation to 1% O_2_ for 24 h (shaded areas), mostly in those cells previously maintained in physioxia.

**Figure 6:**
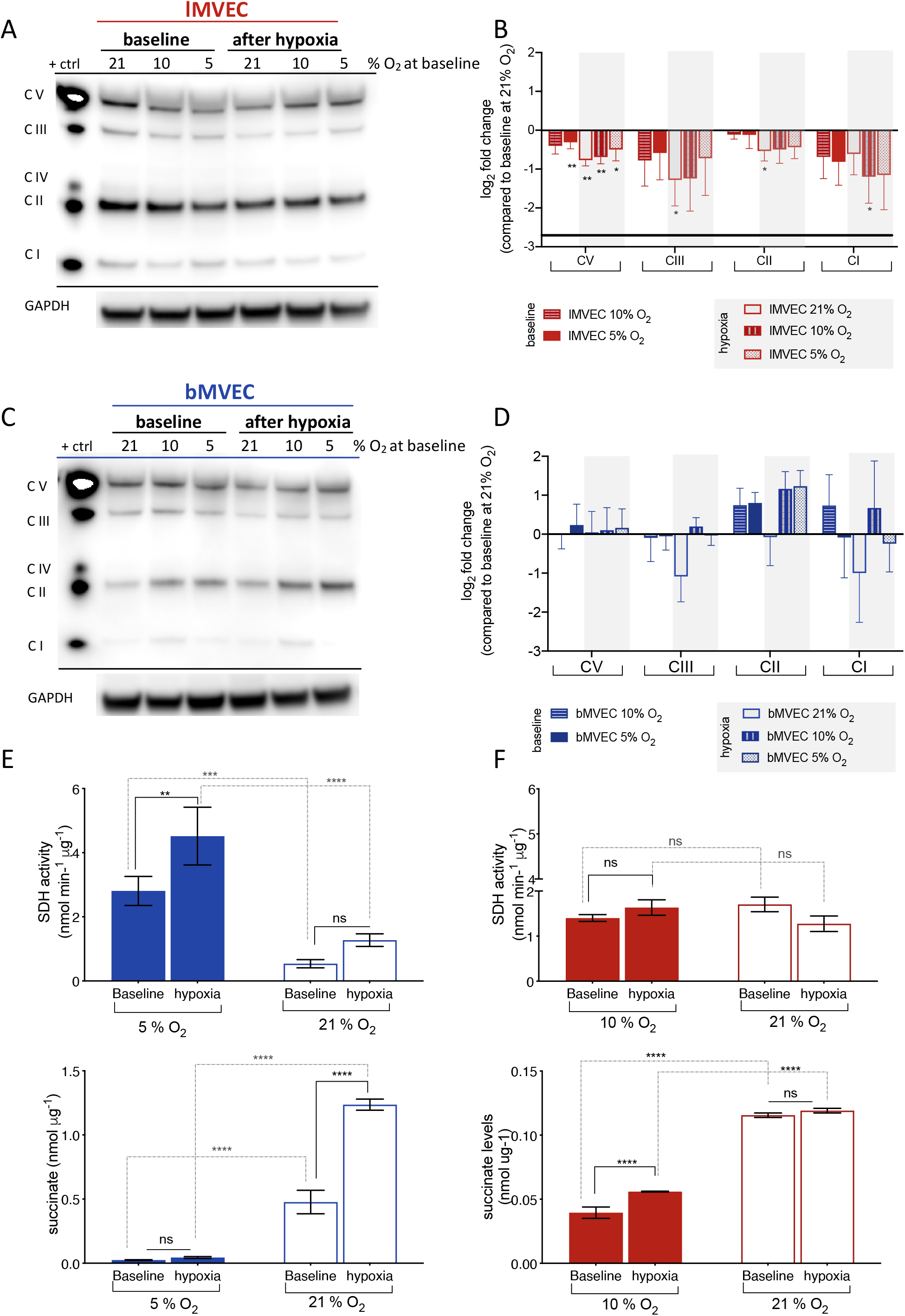
Brain and Lung MVEC have different patterns of mitochondrial ETC components. Whole protein extracts were collected from MVEC from the three baseline (21 %, 10% O_2_ and 5% O_2_) and after exposure of each to 1% O_2_ for 24 h. The lysates were probed by western blot using a mitochondrial antibody cocktail. (A) Representative images are shown for lung MVEC from each O_2_ environment (B) Quantification of the bMVEC western blots was done following densitometry and normalization loading control and each condition compared to levels at 21 % O_2_; Data is shown as log_2_(fold change) ± SD and significance was assessed by one sample t-test; *p<0.05, **p<0.01, n = 4. (C) Representative images are shown for brain MVEC from each O_2_ environment (D) Quantification of the bMVEC western blots was done following densitometry and normalization loading control and each condition compared to levels at 21 % O_2_; Data is shown as log_2_(fold change) ± SD and significance assessed by one sample t-test; *p<0.05, **p<0.01, n = 4. Complex I was detected at levels below threshold for accurate quantification. (E) SDH activity and (F) succinate levels at baseline and after hypoxia (24 h at 1 % O_2_), following priming in physioxia (5 % O_2_ for brain, 10 % O_2_ for lung) and hyperoxia (21 % O_2_). Data is shown as average ± SEM and significance was assessed by 2-way ANOVA, Holm-Sidak’s multiple comparison’s test, ** p<0.01, ***p<0.0001, n=4 for bMVEC, and lMVEC at 10% O_2_, n=2 for lMVEC 21% O_2_

A very different pattern was seen in MVEC that originated from brain tissue (Figure 6C); instead of a general decrease in protein from all ETC complexes, bMVEC cultured at 10 % or 5 % O_2_ have much higher levels of Complex II, which increased following hypoxia exposure (Figure 6C,D). Notably, if MVEC were kept in hyperoxia (21 % O_2_), complex II levels were not changed by hypoxia (Figure 6D). The activity of succinate dehydrogenase (SDH, complex II) (Figure 6E) as well as succinate levels (Figure 6F) were quantified at baseline and following adaptation to hypoxia, in MVEC primed in their own physiological oxygen and compared to those maintained in normal atmosphere. In bMVEC, SDH activity correlated with SDH protein levels shown in Figure 6C,D, and a striking and inverse correlation with succinate levels, which accumulate to significantly higher concentrations in cells from 21 % O_2_, both at baseline and after hypoxia. Physioxic (but not hyperoxic) lMVEC, SDH activity is not affected by culture conditions or hypoxia, but succinate levels (Figure 6F) slightly increase following hypoxia. Interestingly, both baseline levels and the extent of accumulation are an order of magnitude lower than what is seen in bMVEC (note split scales). These data suggest that mitochondria from hyperoxic bMVEC lose SDH protein and activity, and as a result accumulate extremely high (and potentially toxic) levels of succinate. Basal SDH activity and changes in succinate levels in lMVEC change within a much lower scale, indicating that this is a finely tuned but likely less relevant aspect of mitochondrial function MVEC originally from the lung.

The signal intensity for all other quantifiable complexes was not different between any conditions in bMVEC, with the notable exception of much lower levels of complexes III and I in hypoxic cells that were primed at 21 % O_2_. These two complexes are in fact the two mostly associated with ROS generation downstream of mitochondrial activity.

Quantification of mitochondrial potential using MitoTracker Red CMXRos, in which higher signal intensity correlates with mitochondrial health and activity (Supplementary Figure S5), indeed suggested that bMVEC have lower mitochondrial activity at 21 % O_2_, and qualitative observation indicates that hyperoxic mitochondria in lMVEC are more active than those from bMVEC. This could be a result of oxidative damage, to which bMVEC mitochondria appear considerably more susceptible.

These opposing oxygen-induced changes in ETC composition (Figure 6) and mitochondrial activity (Supplementary Figure S5) between the two MVEC populations are consistent with the measurements of maximal mitochondrial respiration shown above (Figure 5), where in bMVEC higher maximal OCR occurred at lower baseline oxygen levels (Figure 5A, B, right), and was higher than that of lMVEC after adaptation to hypoxia, if cells were maintained in physiological oxygen. Conversely, when lMVEC were maintained in physiological O_2,_ they lost their mitochondrial spare capacity almost entirely (Figure 5 C,D, middle and right panels).

## Discussion

MVEC are key regulators of organ homeostasis, and control perfusion and permeability locally to match organ demand (Augustin and Koh, 2017; Reiterer and Branco, 2019). Perception and response to oscillations in oxygen availability, whether as a result of changes in tissue need or environmental supply, are essential aspects of MVEC function (Bartoszewski et al., 2019). In this study, primary MVEC from two continuous endothelial capillary networks were isolated from brain and lung tissue, and their responses to hypoxia compared.

The distinct responses seen here demonstrate that EC are not only intrinsically heterogeneous (Marcu et al., 2018; Nolan et al., 2013; Ribatti et al., 2002), but functionally and metabolically reprogrammable by O_2_ exposure.

The canonical transcriptional hypoxia response is known to rely on activation of the HIF pathway and the stabilization of either or both of the main isoforms of HIF-α. Brain and lMVECs differentially express HIF isoforms, with lMVECs consistently stabilizing higher levels of HIF-1α both at baseline physioxia (but not hyperoxia) and after hypoxia exposure; while HIF-1α is usually associated with acute hypoxia response (Koh and Powis, 2012) the preferred isoform stabilized in bMVEC is HIF-2α, induced by hypoxia to a much higher degree than that seen in lMVEC. Regardless of the preferred HIF-α isoform stabilised in each cell type, however, the mRNA levels of HIF transcriptional targets, specifically those encoding typical hypoxia response genes such as PGK, GLUT1, LDH-A and VEGF, were induced to comparable levels in both MVEC populations (Figure 2), and ultimately these results indicate that the distinct vulnerability to hypoxic insult in these two cell populations is largely HIF-independent.

Endothelial function is intrinsically linked to endothelial metabolism (De Bock et al., 2013; Doddaballapur et al., 2015; Rohlenova et al., 2018; Wong et al., 2017). In this study, baseline metabolic preferences and hypoxia-driven metabolic shifts were measured in the two MVEC and compared to investigate if their distinct responses and tolerance resulted from intrinsic heterogeneity or oxygen priming.

As expected, baseline glycolytic activity was higher MVEC expanded in lower O_2_. Both MVEC grown in physiological atmospheres (5 or 10 % O_2_) had increased maximal glycolytic rates (Figure 3), which correlated with increased viability during hypoxia (Figure 2). As hyperoxygenation impaired the ability to upregulate glycolysis during a subsequent hypoxic stimulus, even in the absence of mitochondrial respiration (Figure 3), the differences in glycolytic capacity appear to motivate loss of cell viability. The timing and magnitude of the metabolic shift upon exposure to hypoxia is not only highly dependent on O_2_ priming, but inherent to the origin of the MVEC (Figure 4). In spite of a relatively narrow challenge (from 5 % to 1 % O_2_), bMVEC are strikingly responsive when compared to the same cells expanded in higher O_2_ conditions, and severely compromised when cells are maintained at 21 % O_2_ (Figure 5).

Interestingly, baseline glucose uptake is not affected by environmental O_2_ and is consistently higher in bMVEC than lMVEC, reflecting the role of brain microvasculature in the active shuttling of glucose to neural tissue (Duelli and Kuschinsky, 2001; Huang et al., 2012; Veys et al., 2020); however, the hypoxia-induced increase in glucose uptake in MVEC from the brain is severely delayed if cells are maintained in supra-physiological O_2_ levels. This had been previously shown in a study on the effect of hyperbaric oxygen on the brain vasculature (such as experienced during diving) (Wilson and Matschinsky, 2019). Combined, these results suggest that hyperoxic priming compromises the ability of bMVEC to perceive the hypoxic challenge, apparently more so than their capacity to respond.

While all EC are assumed to be primarily glycolytic, bMVEC are known to have significantly higher mitochondrial density (Oldendorf et al., 1977), and thus presumably have higher mitochondrial activity. Remarkably, bMVEC only show significantly higher relative mitochondrial respiration rates than lMVEC when expanded in their own physiological atmospheres (5 % O_2_). The dramatic lower maximal respiration of bMVEC relative to lMVEC, when expanded at 21 % O_2_, show that hyperoxia is much more damaging to bMVEC. These are also the O_2_ priming conditions at which bMVEC show a severe delay in induction of glycolysis following hypoxia (Figure 5) and subsequent compromised viability (Figures 1,2).

Mitochondria are obvious sensors of O_2_ availability (Nanadikar et al., 2019; Rabinovitch et al., 2017; Smith and Schumacker, 2019), and even though their cellular function is commonly associated with efficient generation of cellular ATP, in that process they become main generators of reactive oxygen species (ROS)(Dröse, 2013; Larosa and Remacle, 2018; Zhao et al., 2019); these have been shown to mediate mitochondrial damage and associated high rates of cell death in bovine aortic EC exposed to high O_2_ (Wang et al., 2015). Importantly, mitochondrial function has also been associated with the integrity of the BBB (Bukeirat et al., 2016; Doll et al., 2015).

Mitochondrial ROS are generated primarily at complexes I and III of the mitochondrial electron transport chain (mETC) (Larosa and Remacle, 2018; Zhao et al., 2019). Whereas in lMVEC all mETC protein complexes decrease almost linearly with decreased O_2_ availability, the only two complexes seen to decrease in hypoxic bMVEC are precisely complexes I and III, but only if the cells are primed in hyperoxia (Figure 6). Conversely, levels of complex II are higher in physiologically maintained bMVEC, and increase following adaptation to hypoxia, again only if cells are never hyperoxygenated. This suggests that complex II activity is essential for the response in bMVEC to hypoxia, possibly in regulating succinate levels, which are detected at levels 10-fold higher than in lMEVC. Both SDH activity and succinate levels are dramatically affected by hyperoxia in bMVEC. Albeit unclear what the consequences of high succinate might be in these cells, it has been associated with epigenetic regulation of gene expression, and a molecular mediator of signal transduction and metabolic reprogramming (Mills et al., 2016; Tretter et al., 2016). Complex II is a unique transporter in the mETC, the only one that is exclusively encoded in the nuclear genome, and that does not contribute to the proton gradient across the inner mitochondrial membrane (Dröse, 2013). Its role in EC has not been studied, and there is some ambiguity in the existing literature regarding the contribution of this complex to ROS formation (Baysal, 2000; Dröse, 2013; Pfleger et al., 2015; Zhao et al., 2019). However, such studies were performed in hyperoxygenated cells, where such changes would not have been observed (Figure 6D), and to the best of our knowledge, this is the first instance where such assessments were performed while maintaining relevant oxygen tensions. The underlying significance of high succinate levels and high activity of mitochondrial complex II in bMVEC is, however, unclear and worth further investigation, but our data indicates that supra-physiological levels of O_2_ affect the stability of complex II in bMVEC, presumably corrupting reserve respiratory capacity (Figure 5) (Pfleger et al., 2015), and impairing ROS removal. This is supported by the fact that hyperoxygenated bMVEC become unresponsive to hypoxia. If on one hand, having more mitochondria could make cells more sensitive to O_2_ oscillations (Figure 4, right panel), on the other hand cells with more mitochondria will generate more ROS, in direct correlation with O_2_ availability. Thus, the subsequent damage to the mitochondria will in turn compromise the ability to perceive O_2_ oscillations. This too is supported by the fact that bMVECs cultured at 21 % O_2_ fare far worse than those from lung; not only do bMVEC undergo a more severe hyperoxic challenge relative to their physioxic setpoint, but these cells also have more mitochondria (Oldendorf et al., 1977) and higher baseline mitochondrial activity (Figure 5, Supplementary figure S5). While a similar effect can be seen in lMVEC, it is much less severe. As lMVEC are exposed to wider oscillations of O_2_ due to their close proximity to air, they should be inherently more tolerant to higher O_2_ levels and oxidative stress. Most studies regarding the effects of oxygen in microvascular cells, specifically hyperoxia, are restricted to the lung (Ahmad et al., 2004; Ma et al., 2018; Mach et al., 2011; Narula et al., 1998), and occasionally arterial EC (Li et al., 2018; Suzuki et al., 1997; Wang et al., 2015), when microvascular networks from less oxygenated tissues are likely to be much more susceptible.

During hyperbaric oxygen treatments (HOT), used to treat refractory diabetic lower extremity wounds or radiation injuries, O_2_ partial pressure can be up to 15 times higher than standard air (Chen et al., 2015; Goldman, 2009), resulting in a 13-fold increase in oxygenation in rat brain tissue (Thom, 1989). Visible acute side effects (seizures) affect very few patients (Banham, 2011), but early studies in rodents reported a 50 % decrease in lung endothelium viability following exposure to pure O_2_ (Kistler et al., 1967). Even though effects on viability are less pronounced in primates, functional aspects have not been investigated (Kapanci et al., 1969). As of the unfolding the severe acute coronavirus syndrome 2 (SARS-CoV-2) in 2019/2020 pandemic, the use of supplemental oxygen has been also used to combat the resulting hypoxemia, but there is no clear consensus about what the target blood oxygen saturation should be; suggestions range from 90% up to >96%(Ni et al., 2019; Rasmussen et al., 2018; Shenoy et al., 2020), with most treatment guidelines opting for intermediate values; although the discussion hinges on patient mortality as the main indicator, if no differences in mortality are observed, our data suggests that liberal use of oxygen should be met with caution. Previous data showed that perioperative hyperoxia increases mortality in patients with pre-existing pathologies (Damiani et al., 2014; Helmerhorst et al., 2015; Wijesinghe et al., 2008), and so the benefits of this practice should also be reassessed (Mattishent et al., 2019; Volk et al., 2017).

This study highlights the detrimental effects of hyperoxia, which are far less considered although demonstrably no less harmful, and underscores the importance of research into organ-specific MVEC physiological and metabolic reprogramming and downstream consequences in disease, treatment, and long-term effects following environmental challenges.

## Acknowledgements

This work was funded by CRUK Cambridge Centre non-clinical training award to CB/MR (ref. C9685/A23214), and a generous start-up package from the School of Medicine, Dentistry and Biomedical Sciences, Queen’s University, Belfast. The authors would like to thank the support from colleagues and peers from Medina lab at WWIEM (CEM-QUB) for insight and technical support with metabolic analyses.

## Author Contributions

Conceptualization, CBP, RSJ. Methodology, MR, RSJ, CBP. Investigation, MR, AJE, AB, CB; Writing – Original Draft, CBP. Writing – Review & Editing, CBP, RSJ, MR; Funding Acquisition, CBP; Resources, RSJ and CBP; Supervision, RSJ, CB.

## STAR Methods

### Isolation of brain MVECs

Brain MVEC were isolated by incorporating and slightly modifying previously described methods (Coisne et al., 2005; Ruck et al., 2014; Watson et al., 2013; Welser-Alves et al., 2014). Brains of 6-8 wk old male C57BL6/J mice (4-6 animals per isolate), were excised and stored in serum-free DMEM on ice before surgical removal of the olfactory bulbs, cerebellum, and mid-brain white matter. The remaining cortical tissue was rolled on sterile filter paper to remove outer vessels and meninges, and tissue was digested in DMEM containing 2 mg/mL collagenase A (Roche) and 10 µg/mL DNase I (Roche) at 37 °C for 1 h, with gentle rotation. Digested tissue was pelleted at 290 *x g* and resuspended in DMEM containing 20 % BSA (w/v), centrifuged for 10 min at 1000 *x g* to separate the vessel pellet from the buoyant myelin fraction. The cell pellet was resuspended in DMEM and filtered through a 70 µM nylon mesh, to remove large vessel fragments, and the filtrate was centrifuged at 290 *x g*. Pellet was resuspended in a digestion mix containing 2 mg/mL collagenase/dispase (Roche) and 10 µg/mL DNase I in DMEM and incubated on a shaker at 37 °C for 30 min. Following centrifugation at 290 *x g*, microvascular cell pellets were washed once in DMEM, resuspended in MVEC growth medium supplemented with 4 µg/mL puromycin (Sigma) and plated on cell culture plates precoated with collagen I. Puromycin was maintained in the culture medium for the first 4 days to inhibit growth of non-endothelial cells.

### Isolation of lung MVEC

Lung MVEC were isolated as described previously (Branco-Price *et al*., 2012) with slight modifications. The same mice used for brain tissue were used, and lungs excised into DMEM, kept on ice. Finely minced lung tissue was digested for 90 min at 37 °C in HBSS containing collagenase A (2 mg/ml), supplemented with 2 mM CaCl_2_, 2 mM MgSO_4_, and 20 mM HEPES. Digestion suspension was filtered through a 70 µM nylon mesh and washed once in HBSS. Cell pellet was resuspended in PBS containing 0.1% BSA and incubated with anti-rat IgG Dynabeads (Invitrogen) bound to rat anti-mouse CD31 antibody (BD Pharmingen) for 90 min at 4 °C. The beads were washed three times with 0.1 % BSA in PBS, resuspended in MVEC growth medium, and plated on cell culture plates precoated with collagen I.

### MVEC culture and hypoxia treatment

TC plates were coated with collagen (3 mg/mL, Sigma, diluted 1:10 in H_2_O, and incubated for 2 h at 37 °C). Excess solution was removed and wells rinsed twice with PBS immediately prior to plating. MVEC were expanded in a 1:1 mixture of low-glucose DMEM (Gibco) and F12 HAM nutrient mixture (Sigma), buffered with 20 mM HEPES (Gibco), and supplemented with 1 % nonessential amino acids (Sigma), 2 mM sodium pyruvate (Gibco), 20 % FBS (Gibco), 75 µg/mL endothelial cell growth supplement (Sigma), and 100 µg/mL heparin (Sigma), and in atmospheres containing 5 % CO_2_ and either ambient (∼21 %) or physiological O_2_ – corresponding to *in vivo* levels in lung (10 %) and brain (5 %)(Dings et al., 1998; Miller et al., 2010; Wild et al., 2005). All media and solutions were pre-equilibrated with the appropriate oxygen concentration 12 h prior to media changes, trypsinization or cell treatments. All hypoxia experiments were carried out in an atmosphere containing 5% CO_2_ and 1% O_2_ at 37 °C, with controlled humidity, using either a Ruskinn Sci-Tive or a Whitley H35 HEPA hypoxystation. Cells received fresh and pre-equilibrated for 24 h at 1% O_2_ before hypoxia treatments (Newby, Marks and Lyall, 2005).

### Viability time course using Propidium Iodide

Brain and lung MVEC were grown to 90% confluence before being transferred to 1% O_2_. At each timepoint, cells were washed once in PBS, detached using 0.25% trypsin, and resuspended in fresh media. All reagents used had been pre-equilibrated to 1% O_2_. Cell viability was assessed using an Adam-MC automated cell counter (NanoEnTek) according to manufacturer’s instructions. Cells were incubated with either a total cell stain, containing Propidium Iodide (PI) and a lysis agent, or with a non-viable cell stain, containing only PI. Viability was measured as a percentage of non-viable cells compared to total cell number.

### Real time viability

Real time viability was assessed using the RealTime-Glo™ MT Cell Viability Assay (Promega) according to manufacturer’s instructions. 3000 cells were plated one day before the start of the assay on collagen-coated white 96-well plates. Immediately before the start of the assay, MT Cell Viability Substrate and NanoLuc enzyme were added to growth medium pre-equilibrated to the appropriate O_2_. Luminescence was read every 10 min over 48 h, using a FLUOstar Omega plate reader set to 37 °C, 5% CO_2_, and at the appropriate oxygen level using an Atmospheric Control Unit (BMG Labtech).

### qPCR

Brain and lung MVEC were grown to 90% confluence, and either transferred into a hypoxia chamber containing 1% O_2_, or maintained at the same oxygen concentration. RNA was isolated using the RNeasy isolation kit (Qiagen) according to manufacturer’s instructions. cDNA was synthesised from 1 µg of RNA using SuperScript III reverse transcriptase (Invitrogen) according to manufacturer’s instructions. All transcript levels were measured in triplicates and normalised to b-actin; fold-changes were calculated in relation to the 4 h baseline reading.

### Western blot analysis

Brain and lung MVEC were grown to 90% confluence. The cells were then either transferred into a hypoxia chamber containing 1% O_2_ or maintained at the same oxygen concentration in which they were cultured since isolation. In both cases, a media change was carried out at t = 0 using oxygen-equilibrated growth medium. Cytoplasmic and nuclear protein were collected after the indicated times using NE-PER Nuclear and Cytoplasmic Extraction Reagents (Thermo-Scientific) according to manufacturer’s instructions. Total protein was extracted with RIPA buffer containing protease inhibitors (Roche). Protein concentration was measured using a Pierce™ BCA Protein Assay Kit (Thermo Scientific). Unless specified otherwise, the amount of protein used per lane was 3 µg for nuclear protein and 20 µg for cytoplasmic and total protein. Samples were resolved in 3-8% Tris-Acetate gels (Invitrogen) or 4-12% Bis-Tris gels (Invitrogen), and subsequently transferred to PVDF membranes using semi-dry blotting cassettes (Power Blotter XL, ThermoFisher Scientific or Trans-Blot Turbo, Bio-Rad). The membranes were probed with primary antibodies o/n at 4 °C followed with HRP-conjugated secondary antibodies for 1 h at room temperature. All antibodies were diluted in PBST containing 2 % milk. Target bands were detected with Pierce ECL western blot substrate (Thermo Scientific), and quantification was performed using ImageJ.

### Western blot for mitochondrial electron transfer chain complexes

90% confluent cells were transferred to 1% O_2_ or maintained at the same baseline O_2_ concentration for 24 h. Protein was collected with ice-cold RIPA buffer and quantified using the Pierce™ BCA Protein Assay Kit (ThermoFisher). 35 µg of protein per sample were separated on 4-12% Bis-Tris gels and transferred to PVDF membranes. As per manufacturer’s advice, samples were not heated prior to gel-separation to prevent damage to mitochondrial complex I, unstable above 50 °C. Total OXPHOS Rodent WB Antibody Cocktail (Abcam) was diluted 1:250 in PBST containing 1% milk. Western blot steps were performed as above. Bands were visualized using Amersham ECL Western Blotting Detection Reagent (GE Healthcare). Rat heart mitochondrial extract was used as a positive control.

### Glucose uptake assays

3 × 10^4^ MVEC were seeded per well in a 96-well plate 12 h before the start of the assay, and either transferred into a hypoxia chamber containing 1% O_2_ or maintained at the same oxygen concentration. The start of each hypoxia incubation was staggered such that 2-DG treatment could be carried out at the same time for all conditions. The cells were washed once with PBS and incubated with 0.1 mM 2-DG in PBS for 10 min; glucose uptake was measured using the Glucose Uptake-Glo™ Assay (Promega) according to manufacturer’s instructions. All reagents used were pre-equilibrated to the appropriate O_2_. Samples were incubated with detection reagent for 2 h and luminescence was measured with a FLUOstar Omega plate reader. A standard curve of 2-deoxy-glucose-6-phosphate was used to calculate glucose uptake rates. Replicate wells plated at the same time were used to determine the cell number at the time of 2-DG treatment.

### Glycolytic and mitochondrial stress tests

Local pH and O_2_ changes in media were measured using a XFe96 Analyzer (Agilent). Appropriate cell density was optimised for each measurement and condition, as mitochondrial stress tests required a lower cell number to avoid anoxia in the well after FCCP treatment; 8,000 MVEC were used for mitochondrial stress tests at baseline, and 5,000 at 1% O_2_; For glycolytic stress tests, 10,000 cells were used. MVEC were plated on Seahorse microplates precoated with collagen I and left to adhere for 12 h, then assayed immediately (baseline readings) or transferred to 1% O_2_ for 24 h before assay. Immediately prior to transfer to hypoxia, all wells received fresh medium pre-equilibrated to 1% O_2_. Before mitochondria stress tests, all wells were washed twice with Seahorse XF base medium supplemented with 7.8 mM glucose, 2.5 mM glutamine, and 3.5 mM pyruvate. For glycolytic stress tests, Seahorse XF base medium was supplemented with 2.5 mM glutamine only. The pH of the medium was adjusted to 7.4 using 1 M NaOH. The cells were then equilibrated with assay medium for 45 min at 37 °C in a CO_2_-free atmosphere at the appropriate O_2_ level. After recording baseline measurements, for mitochondrial stress tests, 1 mM Oligomycin, 1 mM FCCP and combined Antimycin A and Rotenone (0.5 mM each) were added successively. For glycolytic stress tests, successive injections of 10 mM Glucose, 1 mM Oligomycin and 50 mM 2-deoxy glucose were sequentially injected. Changes in pH and oxygen consumption were tracked in real time after each injection(Chang et al., 2018; Li et al., 2017; Rellick et al., 2016). Each measurement was composed of a mix-wait-measure cycle which lasted 5 min – 1 min – 2min (XF24 in hypoxia and normoxia), 5 min – 0 min – 2 min (XFe96 in hypoxia), or 3 min – 0 min – 3min (XFe96 in normoxia), as recommended by the manufacturer. Protein concentration in each well was measured using Pierce™ BCA Protein Assay Kit (ThermoFisher) after assay, and extracellular acidification rates (ECAR) and oxygen consumption rates (OCR) were normalised to µg of total protein. Basal and maximal rates, as well as spare capacity of glycolysis and oxidative phosphorylation were calculated as defined by the manufacturer.

### Real time hypoxia response curves

Real time hypoxia response curves were measured using a Seahorse XF24-3 analyser. 3 × 10^4^ MVEC per well were plated the day before the assay on Seahorse microplates precoated with collagen I. At the start of the assay, MVEC growth medium was replaced with Seahorse XF base medium supplemented with glucose (7.8 mM), pyruvate (3.5 mM), and glutamine (2.5 mM), matching the concentrations found in MVEC growth medium. Seahorse medium was equilibrated to 1% O_2_ for 24 h prior to start of assay, and ECAR was measured every 20 min for 24 h. Each measurement was composed of a mix-wait-measure cycle which lasted 5 min – 1 min – 2 min, followed by a time delay of 12 min. ECAR was normalised to µg of total protein.

### Hypoxia chambers for Seahorse analysers

For experiments at non-atmospheric O_2_ conditions, the XF24-3 Analyzer was placed within a gas flow-controlled Perspex hermetic chamber. The atmosphere was CO_2_-free and O_2_ levels were set to either 10%, 5%, or 1%. During the assay temperature was controlled within the Seahorse analyser; internal air circulation within chamber was maintained by a fan. The XFe96 Analyzer was placed in a Ruskinn hypoxia chamber, set to either 10%, 5%, or 1% O_2_ and 0.1% CO_2_, as the machine could not be set to 0% CO_2_. The chamber was humidity and temperature controlled; internal air circulation was maintained by a fan. In both cases, the instruments as well as all media and calibrant were equilibrated to the hypoxic atmosphere for 12 h prior to the assay. To avoid reoxygenation, cell culture plates were transported from their incubators to the instrument chamber in an air-tight container, and all washes were carried out within the chamber.

### Mitochondrial staining

MVECs were seeded on sterile glass chamber slides precoated with collagen I and grown to confluence. The cells were stained with fresh O_2_-equilibrated media containing 200 nM MitoTracker Red CMXRos (ThermoFisher) for 30 min, or DMSO as a negative control. The wells were washed twice with PBS and the cells were fixed with cold acetone (−20 °C) for 10 min and permeabilised with 0.5% Triton-X100 in PBS for 5 min at room temperature. Blocking was done using 10% donkey serum in PBS for 1 h at room temperature. Primary VE-Cadherin antibody was diluted in blocking solution (1:50, R&D Systems) and the slides were probed over night at 4 °C, followed by secondary antibody (Goat IgG conjugated with AlexaFluor 488) at room temperature for 1 h. Secondary-only control wells were kept in blocking solution instead of primary antibody overnight and then received the same dose of secondary antibody. Slides were mounted using ProLong Diamond Antifade with DAPI (ThermoFisher). All images were obtained with a Leica DM5500 B Fluorescence Microscope and analysed using ImageJ.

### Migration assays

Fully confluent MVEC in a 12-well plate were treated for 2 h with 10 mM Mitomycin C (or equal volume of vehicle control) in adequately O_2_-equilibrated, serum-free growth medium. A vertical scratch was applied to the plate using a P1000 tip to create a gap in the cell monolayer. The wells were washed twice with PBS to remove detached cells and incubated with fresh oxygen-equilibrated growth medium. Wells were imaged at T= 0, 2, 4, 8, 24, and 48 h. The area of the scratch was measured using ImageJ and normalised to the area at t = 0 to assess percent closure.

### Succinate dehydrogenase assay

Extracts and assay were performed using a commercial succinate assay kit (Abcam), with minor modifications. Confluent wells of 6-well plates were rinsed in ice-cold PBS and resuspended in 100 mL of ice-cold SDH Assay Buffer (provided), both of which pre-equilibrated to the adequate O_2_ levels (21%, 10%, 5% or 1% O_2_). Cells were scrapped from wells using cell lifters, then transferred to pre-cooled microfuge tubes and homogenized by pipetting. Clear extracts were used for assay, and conversion of substrate (DCIP, provided) was assessed by following changes in absorbance at 600 nm over time. Measurements in kinetic mode were made continuously for 30 min, and linear range selected to calculate changes of absorbance over time (DOD/ DT). DOD was converted to nmol of DCIP following subtraction of background and against a standard curve. Results were normalized to amount of protein used per well.

### Succinate assay

Extracts and assay were performed using a commercial succinate assay kit (Abcam) with minor modifications. Confluent wells of 6-well plates were rinsed in ice-cold PBS and resuspended in 100 mL of ice-cold succinate assay buffer (provided), both of which pre-equilibrated to the adequate O_2_ levels (21%, 10%, 5% or 1%). Cells were scrapped from wells using cell lifters, then transferred to pre-cooled microfuge tubes and homogenized by pipeting. Cleared supernatant was deproteinated through a 10 kDa filter column. Filtrates (10 mL) were used for assay, and background wells (lacking succinate converter) were ran in parallel for each sample.

Endpoint measurements were performed at 450 nm and quantification following a 30 min incubation at 37 °C. Quantification of succinate was performed against a standard curve, after background correction, and subsequently normalized to amount of protein used in each well (calculated from BCA assay, performed prior to deproteination).

### Statistical Analysis

Data was analysed in Prism Graphpad 8.3, replicate number and statistical test are described for each experiment.

## Key Resources Table

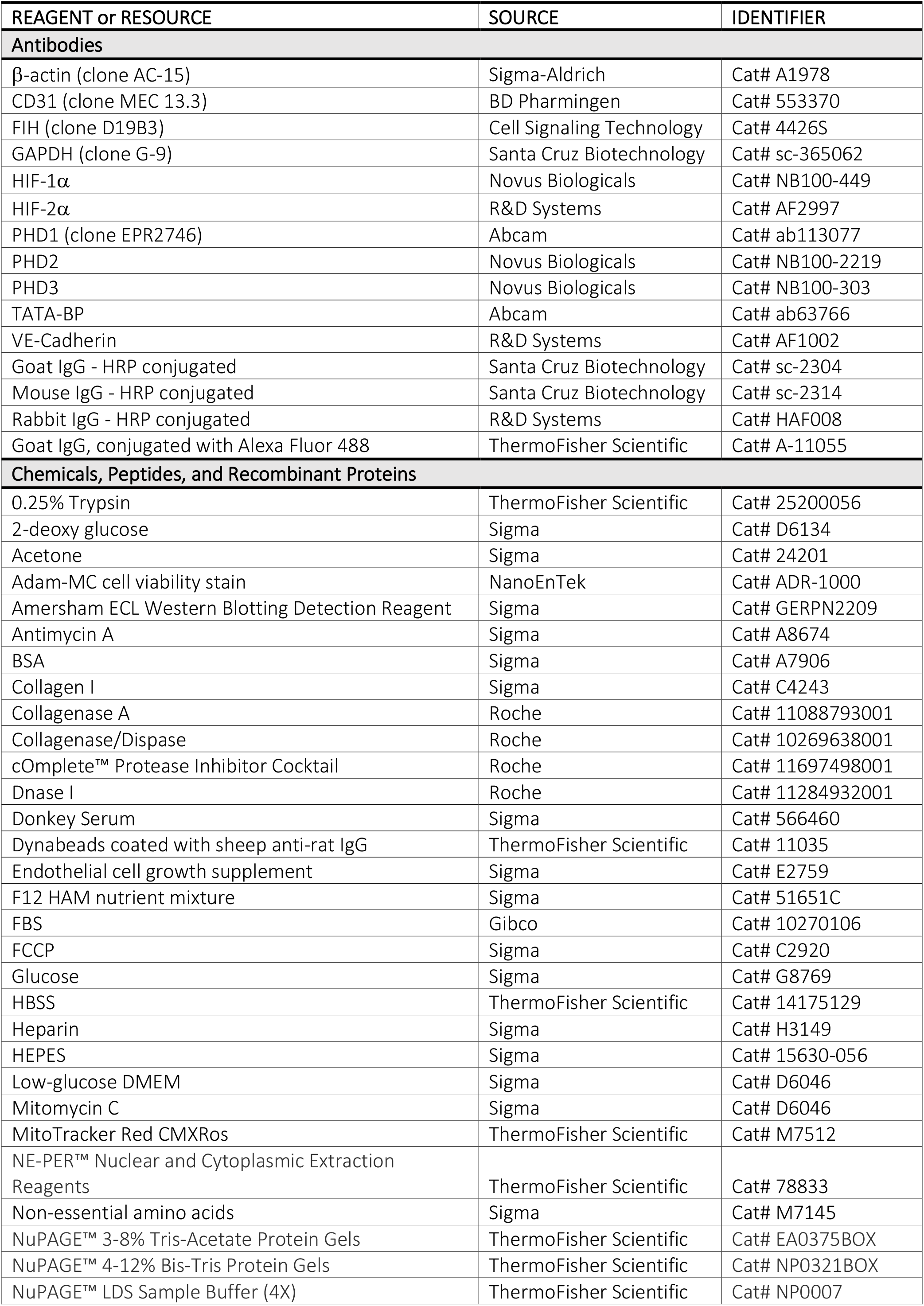

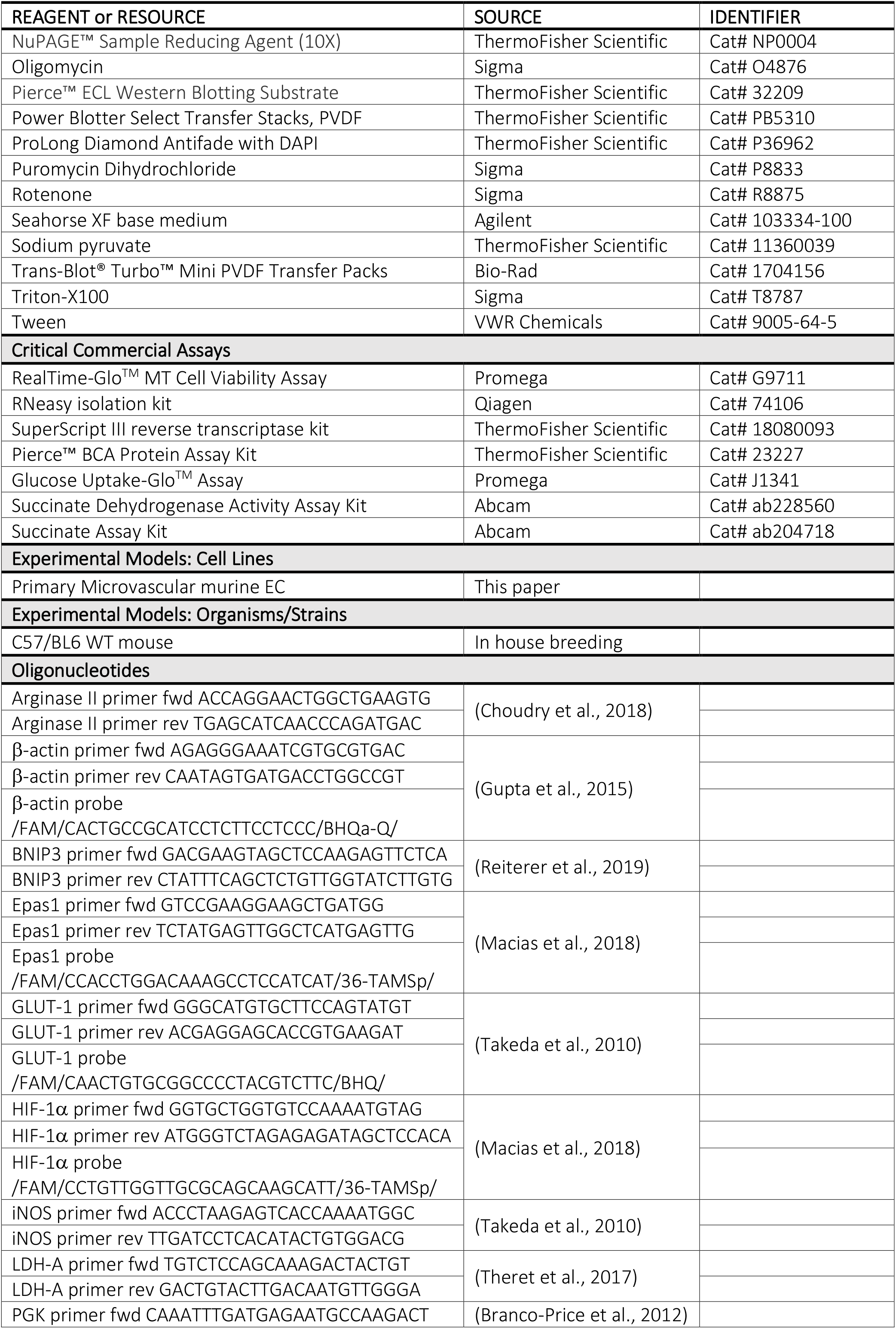

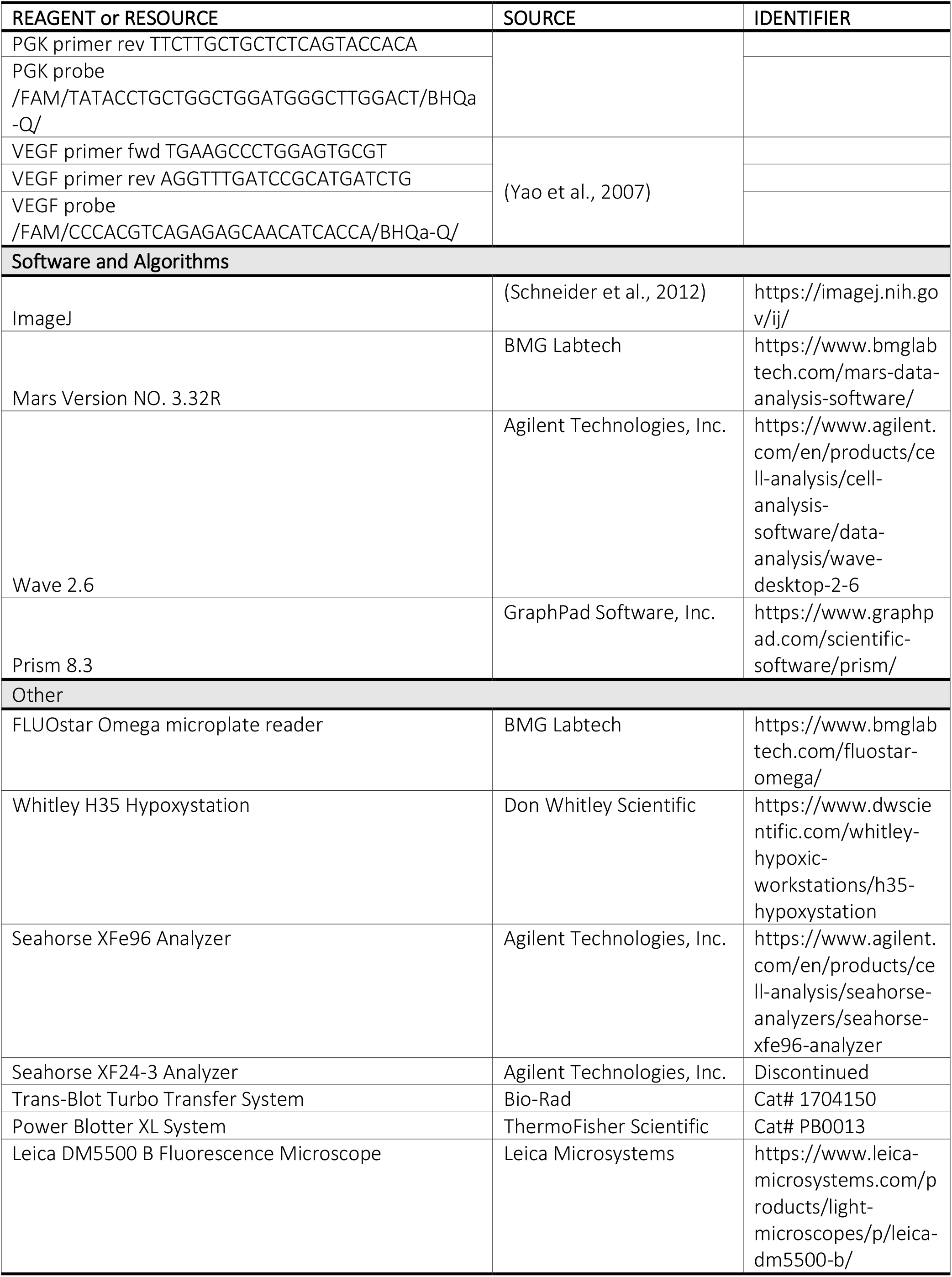

## Resource Availability

### Lead contact

Further information and requests for resources and reagents should be directed to and will be fulfilled by the Lead Contac, Dr Cristina M Branco

### Materials availability

This study did not generate new unique materials.

