## Supplementary Figures for "MICROVASCULAR ENDOTHELIAL CELL ADAPTATION TO HYPOXIA IS ORGAN-SPECIFIC AND CONDITIONED BY ENVIRONMENTAL OXYGEN"

**Figure S1**

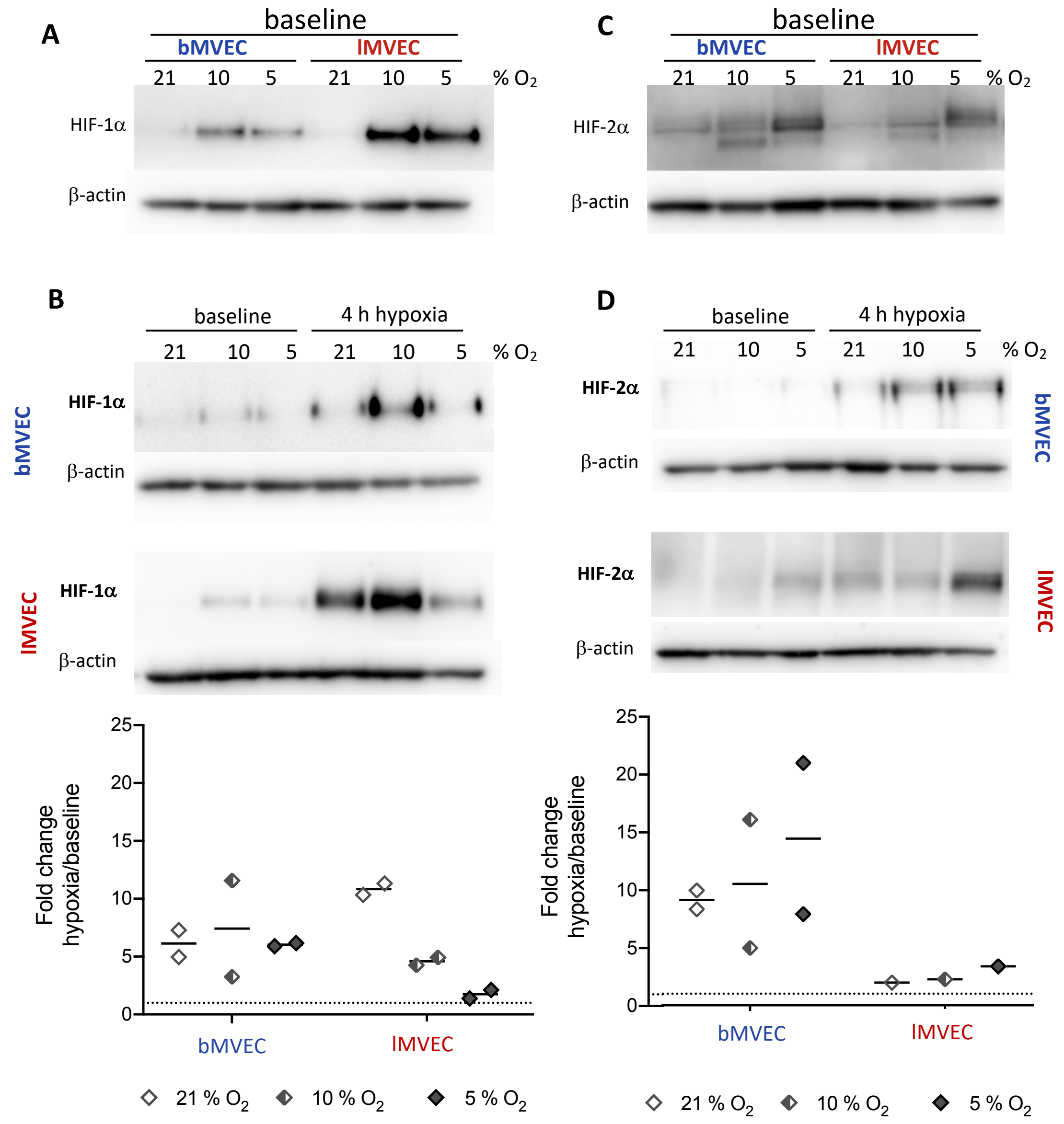


**Figure S1: Baseline and change of HIF-α isoform protein levels is shaped by $O_2$ priming (related to Figure 2)**

(A) Representative western blot imaged showing HIF-1α and HIF-2α signal in nuclear extracts of brain and lung MVEC grown at different oxygen levels; β-actin is shown as loading control

(B) Representative western blot of HIF-1α signal at baseline and following 4h of hypoxia (upper panel), from nuclear extracts of MVEC expanded at different oxygen levels; ratio of hypoxia:baseline signal, normalised to loading control, is shown as fold change (lower panel); n=2

(C) Representative western blot of HIF-2α signal at baseline and following 4h of hypoxia (upper panel), from nuclear extracts of MVEC expanded at different oxygen levels; ratio of hypoxia:baseline signal, normalised to loading control, is shown as fold change (lower panel); n=2

**Figure S2**

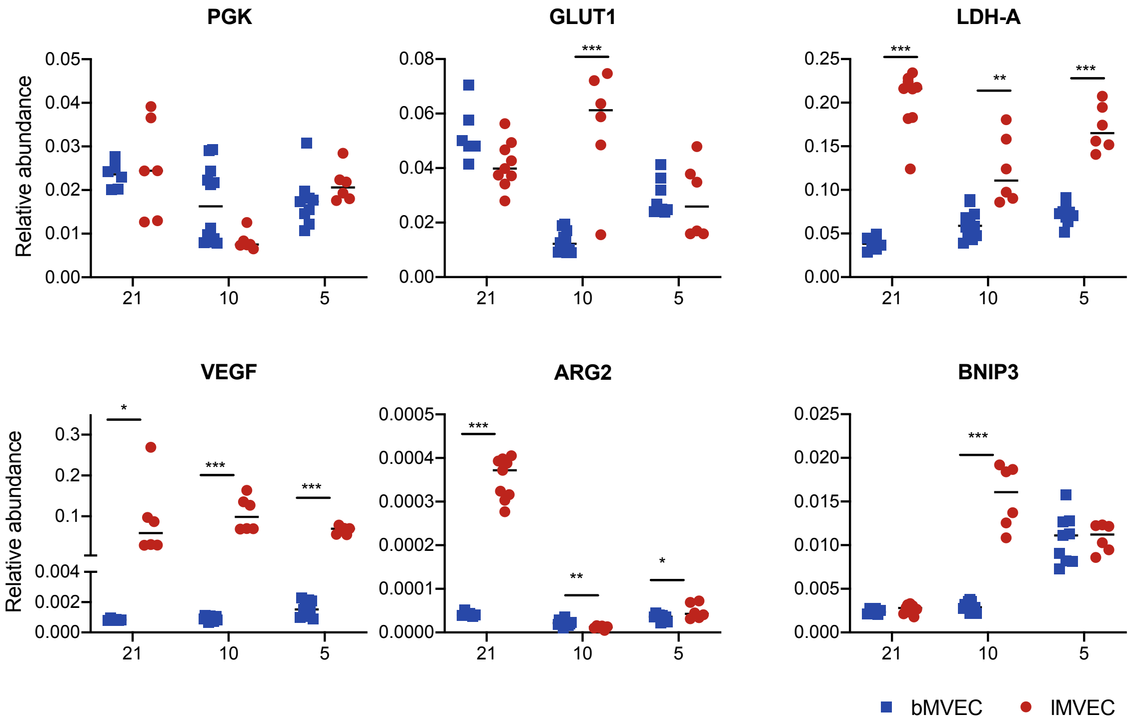


**Figure S2: Relative mRNA abundance of hypoxia response genes under different baseline $O_2$ conditions (Related to Figure 2)**

Total RNA was isolated from brain and lung MVECs expanded at different oxygen levels (21, 10 and 5 % $O_2$). Transcript levels were measured by RT-qPCR and are shown as relative abundance after normalization to the β-actin housekeeping mRNA. n≥2 independent experiments with 3 replicates each; statistical significance was assessed by t-tests corrected for multiple comparisons (Holm-Sidak); *p<0.05, **p<0.01, ***p<0.001

**Table S1: Summary of statistical analyses for figure 2F,** using 2-way ANOVA with Holm-Sidak's multiple comparison test on the log10 of fold change. *p<0.05, **p<0.005, ***p<0.001, ****p<0.0001. When comparing lung versus brain, asterisks are colored to match the higher value, when significant. Note: Arg2 mRNA was undetectable in baseline bMVEC at 21% $O_2$.

| | target | PGK | | | | VEGF | | | | GLUT1 | | | |
|---|---|---|---|---|---|---|---|---|---|---|---|---|---|
| | time | 4h | 8h | 24h | 48h | 4h | 8h | 24h | 48h | 4h | 8h | 24h | 48h |
| | 10 % vs 21 % | **** | **** | **** | **** | **** | **** | **** | **** | ns | **** | **** | **** |
| | 5 % vs 21 % | **** | **** | **** | **** | **** | **** | **** | **** | ns | ns | **** | **** |
| bMVEC | 10 % vs 5% | ns | * | **** | **** | ns | **** | **** | **** | ns | **** | **** | **** |
| | 10 % vs 21 % | ns | ns | **** | **** | ns | ns | ns | **** | ns | * | ns | **** |
| | 5 % vs 21 % | ns | ns | ns | **** | ns | ** | ns | **** | ns | ns | **** | **** |
| lMVEC | 10 % vs 5% | ns | *** | ** | **** | ns | *** | ns | *** | ns | ns | * | ns |
| | 21% | **** | *** | **** | **** | ** | ** | **** | * | ns | ns | **** | **** |
| | 10% | ns | ns | ns | ns | *** | **** | **** | **** | ns | **** | *** | ns |
| lMVEC vs bMVEC | 5% | ns | ** | **** | ** | **** | **** | * | ns | ns | ns | **** | **** |

| | target | ARG2 | | | | LDH | | | | BNIP3 | | | |
|---|---|---|---|---|---|---|---|---|---|---|---|---|---|
| | time | 4h | 8h | 24h | 48h | 4h | 8h | 24h | 48h | 4h | 8h | 24h | 48h |
| | 10 % vs 21 % | ** | **** | **** | - | ns | ns | **** | **** | **** | **** | ns | * |
| | 5 % vs 21 % | ns | * | **** | - | ns | ns | ns | ns | ** | **** | **** | **** |
| bMVEC | 10 % vs 5% | * | **** | **** | *** | ns | ns | **** | **** | ns | **** | **** | **** |
| | 10 % vs 21 % | ns | ** | ns | **** | ** | ns | **** | **** | ns | * | ** | **** |
| | 5 % vs 21 % | ns | ns | ns | **** | ns | * | ns | ns | ns | ns | ** | *** |
| lMVEC | 10 % vs 5% | ns | *** | *** | ns | **** | **** | *** | **** | ns | ns | ns | *** |
| | 21% | ns | ** | **** | - | * | ns | ns | ns | ** | **** | **** | **** |
| | 10% | ns | ns | ns | ns | ns | ns | ns | *** | ns | **** | ns | *** |
| lMVEC vs bMVEC | 5% | ns | ns | ns | ns | ns | * | ns | ns | ns | ns | **** | ns |

**Figure S3**

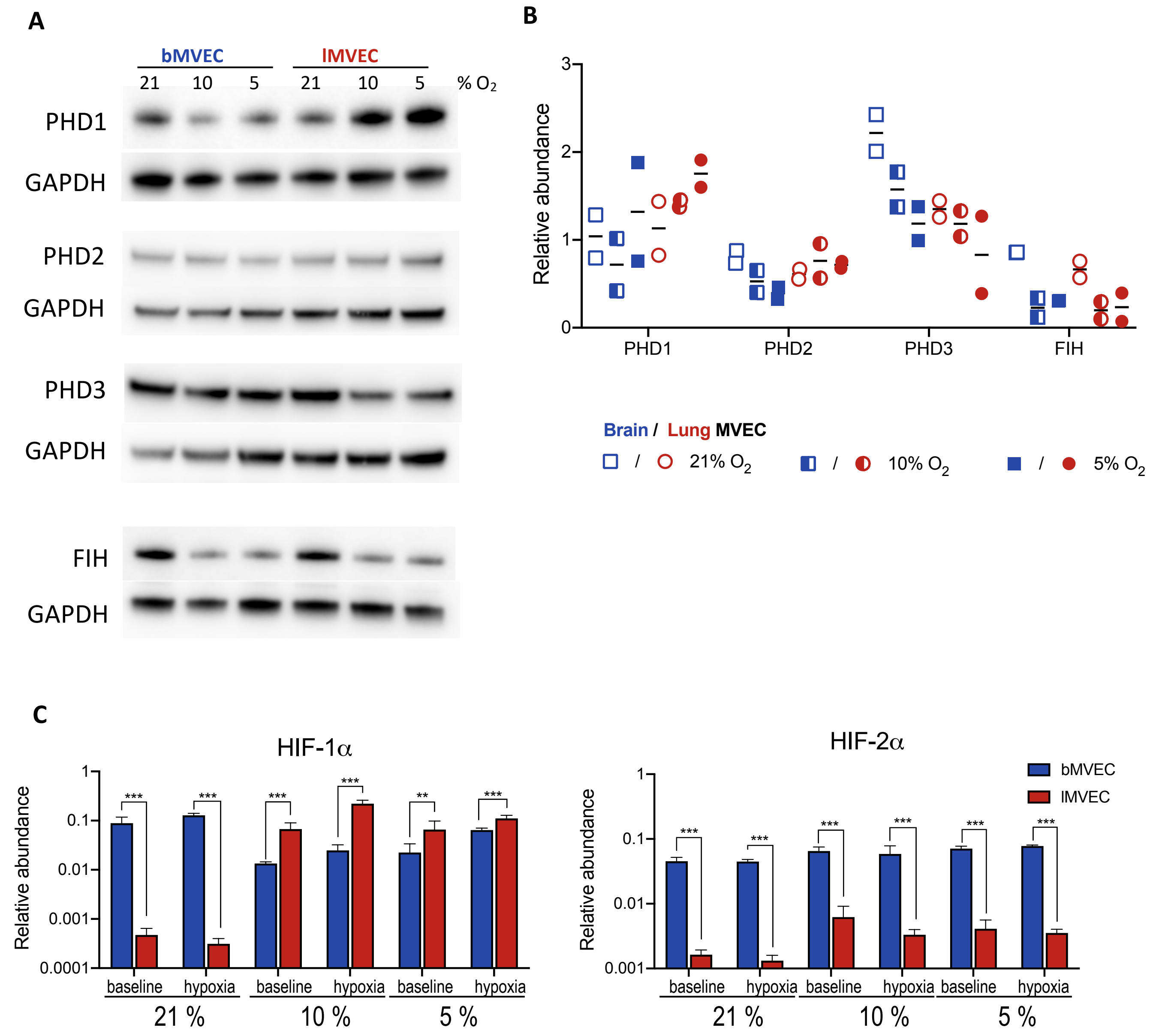


**Figure S3: Upstream regulation of HIF-α is mildly affected by $O_2$ but is not tissue-specific (Related to Figure 2)**

(A) Representative image of western blot probed for enzymes responsible for HIF protein stabilization (prolyl hydroxylases, PHD1-3) and HIF transcriptional activity (factor inhibiting HIF, FIH) using whole-cell protein lysate.
(B) Signal from (A) was quantified by densitometry and normalised to loading control, GAPDH. (n=2)
(C) Comparison of HIF-1α and HIF-2α mRNA levels MVEC from lung and brain, expanded in different oxygen levels, at baseline and after 4h at 1% $O_2$. Transcript levels were quantified by RT-qPCR and are displayed as average ± SD relative abundance compared to the β-actin housekeeping gene (n=3); statistical significance assessed by t-tests corrected for multiple comparisons (Holm-Sidak); **p<0.01, ***p<0.001

**Figure S4**

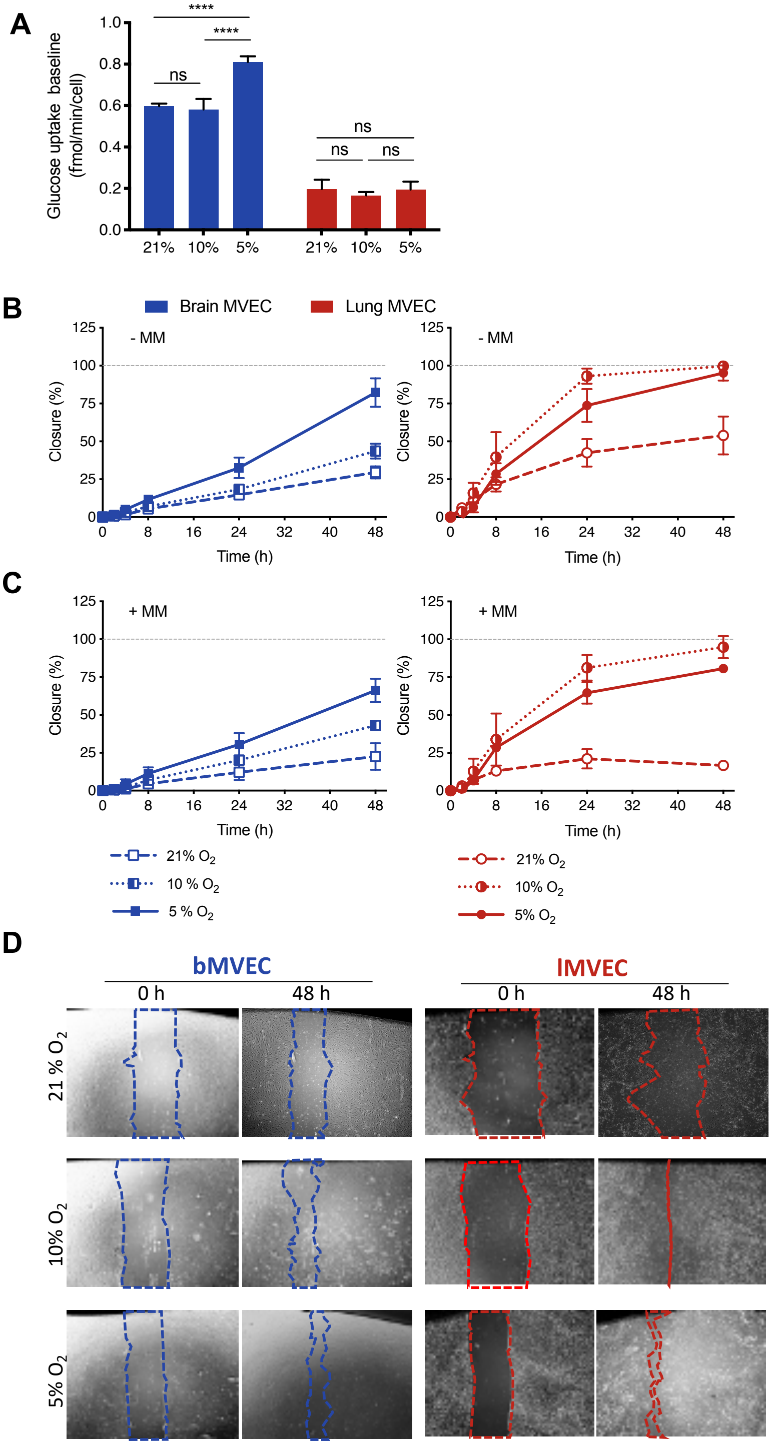


**E**

| - MM | bMVEC | | | | | |
|---|---|---|---|---|---|---|
| time | 0 | 2 | 4 | 8 | 24 | 48 |
| 21 % vs 10% | ns | ns | ns | ns | ns | *** |
| 21% vs 5 % | ns | ns | ns | ns | **** | **** |
| 10 % vs 5 % | ns | ns | ns | ns | *** | **** |

| - MM | lMVEC | | | | | |
|---|---|---|---|---|---|---|
| time | 0 | 2 | 4 | 8 | 24 | 48 |
| 21 % vs 10% | ns | ns | ns | **** | **** | **** |
| 21% vs 5 % | ns | ns | ns | ns | **** | **** |
| 10 % vs 5 % | ns | ns | ns | ** | **** | **** |

| - MM | Lung vs Brain MVEC | | | | | |
|---|---|---|---|---|---|---|
| time | 0 | 2 | 4 | 8 | 24 | 48 |
| 5 % | ns | ns | ns | **** | **** | *** |
| 10 % | ns | ns | *** | **** | **** | **** |
| 21 % | ns | ns | ns | *** | **** | **** |

| + MM | bMVEC | | | | | |
|---|---|---|---|---|---|---|
| time | 0 | 2 | 4 | 8 | 24 | 48 |
| 21 % vs 10% | ns | ns | ns | ns | ns | **** |
| 21% vs 5 % | ns | ns | ns | ns | *** | **** |
| 10 % vs 5 % | ns | ns | ns | ns | ns | **** |

| + MM | lMVEC | | | | | |
|---|---|---|---|---|---|---|
| time | 0 | 2 | 4 | 8 | 24 | 48 |
| 21 % vs 10% | ns | ns | ns | *** | **** | **** |
| 21% vs 5 % | ns | ns | ns | ** | **** | **** |
| 10 % vs 5 % | ns | ns | ns | ns | *** | *** |

| + MM | Lung vs Brain MVEC | | | | | |
|---|---|---|---|---|---|---|
| time | 0 | 2 | 4 | 8 | 24 | 48 |
| 5 % | ns | ns | ns | *** | **** | *** |
| 10 % | ns | ns | ns | **** | **** | **** |
| 21 % | ns | ns | ns | ns | ns | ns |

**Figure 4: Oxygen effects on glucose uptake and cell migration**

(A) Baseline glucose uptake was quantified in lung and brain MVEC maintained in different $O_2$ conditions. Displayed as average ± SD, statistical analysis using 2-way ANOVA with Holm-Sidak's multiple comparison test, ****p<0.0001 (Related to Figure 3)

(B, C) Wound closure assays were performed using brain and lung MVEC expanded at different $O_2$ levels; wounds were applied using a P1000 tip at t=0 and images taken at regular intervals. Quantification of wound closure was made in monolayers treated with DMSO (B) or 10 mM Mitomycin C (C) for 2h before the scratch was applied.

(D) Representative images of wells during migration assay in the presence of Mitomycin C at the start and end of assay.

(E) Summary of statistical analyses for data in B,C; 2-way ANOVA. Multiple comparisons using Holm-Sidak correction; **p<0.005, ***p<0.001, ****p<0.0001, n= 4.

**Figure S5**

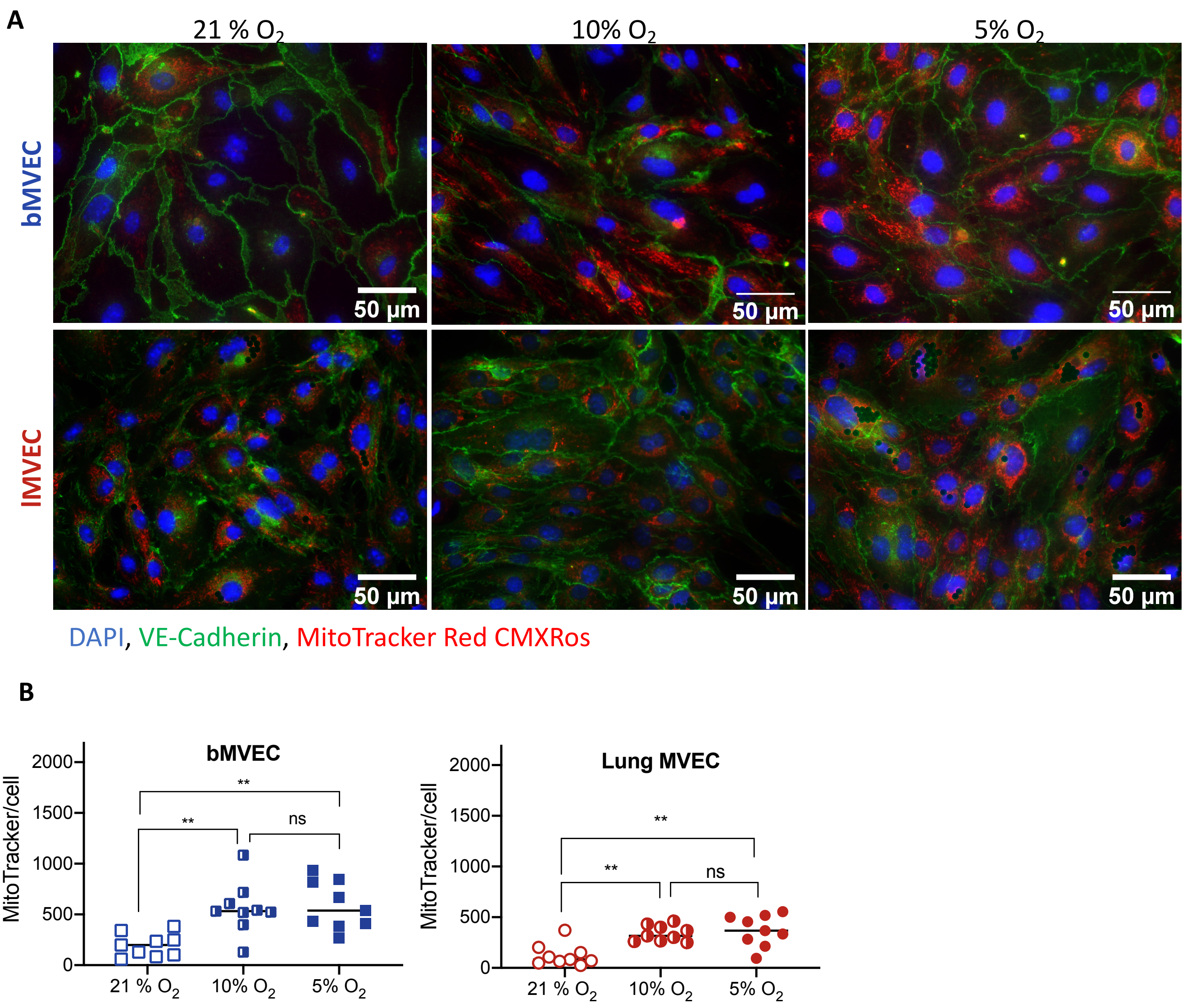


**Figure S5: Mitochondrial activity is impaired by excess $O_2$**

(A) Representative images of brain and lung MVEC expanded at different $O_2$ levels, stained for EC marker VE-Cadherin (green), mitochondrial membrane potential (MitoTracker Red CMXRos, red) and counter-stained with DAPI (blue)

(B) Quantification of the MitoTracker stain shown in (A); different thresholds were used for the two cell populations; n=10 pictures from a single biological control. Statistical analysis using 2-way ANOVA with Holm-Sidak's multiple comparison test (**p<0.005)
